# Genetic architecture of floral traits in bee- and hummingbird-pollinated sister species of *Aquilegia* (columbine)

**DOI:** 10.1101/2021.04.12.439277

**Authors:** Molly B. Edwards, Gary P. T. Choi, Nathan J. Derieg, Ya Min, Angie C. Diana, Scott A. Hodges, L. Mahadevan, Elena M. Kramer, Evangeline S. Ballerini

## Abstract

Interactions with animal pollinators have helped shape the stunning diversity of flower morphologies across the angiosperms. A common evolutionary consequence of these interactions is that some flowers have converged on suites of traits, or pollination syndromes, that attract and reward specific pollinator groups. Determining the genetic basis of these floral pollination syndromes can help us understand the processes that contributed to the diversification of the angiosperms. Here, we characterize the genetic architecture of a bee-to-hummingbird pollination shift in *Aquilegia* (columbine) using QTL mapping of 17 floral traits encompassing color, nectar composition, and organ morphology. In this system, we find that the genetic architectures underlying differences in floral color are quite complex, and we identify several likely candidate genes involved in anthocyanin and carotenoid floral pigmentation. Most morphological and nectar traits also have complex genetic underpinnings; however, one of the key floral morphological phenotypes, nectar spur curvature, is shaped by a single locus of large effect.

## Introduction

Pollinator interactions are a major force in shaping the evolutionary trajectory of flowering plants (Darwin 1878; Crepet 1984; Grimaldi 1999). While angiosperm species can be generalists and attract multiple different pollinators, many have evolved suites of traits, or pollination syndromes, that attract and reward specific pollinators to maximize pollen transfer (Grant 1949; Fenster et al. 2004). For example, the bee pollination syndrome is typified by showy non-red petals, a floral landing platform, and concentrated nectar; hummingbird-pollinated flowers tend to be red, have long tubular corollas and exerted reproductive organs, and provide abundant dilute nectar; and flowers pollinated by hawkmoths have even longer corolla tubes, and produce heady fragrances but little floral pigment to aid in nocturnal detection. Distantly related species have converged on these pollination syndromes: for instance, species in genera as disparate as *Costus* (Costaceae), *Phygelius* (Scrophulariaceae), and *Aquilegia* (Ranunculaceae) all exhibit the suite of floral traits typical of hummingbird-pollination (Cronk and Ojeda 2008). Closely-related groups have also experienced multiple, independent shifts to the same pollination syndrome; *Penstemon* is an extreme example, with an estimated 10-21 independent transitions from bee to hummingbird pollination having occurred within the genus during a recent rapid radiation (Wilson et al. 2007). Indeed, pollinator specialization is associated with species diversity and rapid speciation events in the angiosperms (Armbruster and Muchhala 2009).

The changes in individual floral traits that accompany these pollination syndrome shifts are profound, and must also be coordinated so that the whole flower can reach a new adaptive optimum for pollinator attraction, reward, and pollen transfer. Illuminating the genetic underpinnings of floral trait evolution during pollinator-driven rapid radiation can give us insight into some of the processes that generated the staggering diversity of the angiosperms. A common hypothesis is that biochemical traits such as flower color and nectar composition are controlled by few loci of large effect, such that a change in a single gene can lead to dramatic alteration of phenotype (Rockman 2012; Sheehan et al. 2012). On the other hand, morphological traits such as petal shape and stigma length are generally thought to be governed by many loci of minor effect. Quantitative trait locus (QTL) mapping studies in systems with contrasting pollination syndromes suggest these patterns hold true to some extent, but exceptions are common (Galliot et al. 2006; Hermann and Kuhlemeier 2011). As sequencing technology becomes increasingly accessible and more species are studied, it is unclear how universal this dichotomy between biochemical and morphological traits will remain.

The genus *Aquilegia* (columbine, Ranunculaceae) is an ideal study system for examining the genetic basis of pollination syndrome evolution. It is characterized by a key innovation, the petal nectar spur, that allows for pollinator specialization and facilitated the rapid radiation of the genus (Hodges and Arnold 1995). *Aquilegia* originated approximately 6.9 million years ago in eastern Asia; while there is considerable species diversity in Eurasian taxa, the vast majority exhibit the bee-pollination syndrome, characterized by blue-purple flowers, and petals with long blades for landing as well as short curved nectar spurs (Hodges et al. 2004; Fior et al. 2013). When the genus migrated to North America c. 4.8 million years ago, it experienced rapid diversification to adapt to new pollinators (Whittall and Hodges 2007; Fior et al. 2013). Early in this radiation, there were two independent transitions to hummingbird-pollination, characterized by elongation and straightening of the petal nectar spur, reduction of the petal blade length, exertion of the reproductive organs, and a shift from blue to red floral pigmentation. Subsequently, there were five shifts from hummingbird to hawkmoth pollination, involving further elongation of the nectar spur (up to 16cm in the case of *A. longissima*), loss of anthocyanin pigments, and acquisition of floral scent. There are hawkmoths in Eurasia, but *Aquilegia* only experienced this pollination syndrome evolution once it reached North America, where there are both hummingbirds and hawkmoths. Whittall and Hodges (2007) suggested that hummingbirds, with tongue lengths in between those of bees and moths, served as a necessary intermediate in the evolution of spur length and accompanying floral trait shifts. Therefore, characterizing this initial transition from bee-to hummingbird-pollination is essential for understanding the diversification of the genus as a whole.

In this study, we use QTL mapping of floral traits in the only pair of *Aquilegia* sister species with these contrasting pollination syndromes to begin to elucidate the genetic underpinnings of the genus’s rapid radiation, and to assess whether any patterns are in line with QTL studies of pollination syndromes in other systems. *A. brevistyla* exhibits the bee syndrome, and is found in alpine regions in the northwestern United States and Canada (Fig. 1A; Roe 1992). Its sister species, *A. canadensis*, is hummingbird-pollinated and has a more cosmopolitan distribution across much of North America (Fig. 1A; Herlihy and Eckert 2005). They represent the only pair of sister species in the genus with bee- and hummingbird-pollination, and are estimated to have diverged less than 3 million years ago (Whittall and Hodges 2007; Fior et al. 2013). This study is the first genetic characterization of a bee-to-hummingbird shift in *Aquilegia*, and is unique in the breadth and depth of traits examined, with 17 floral traits spanning color, nectar composition, and organ morphology.

**Figure 1.**
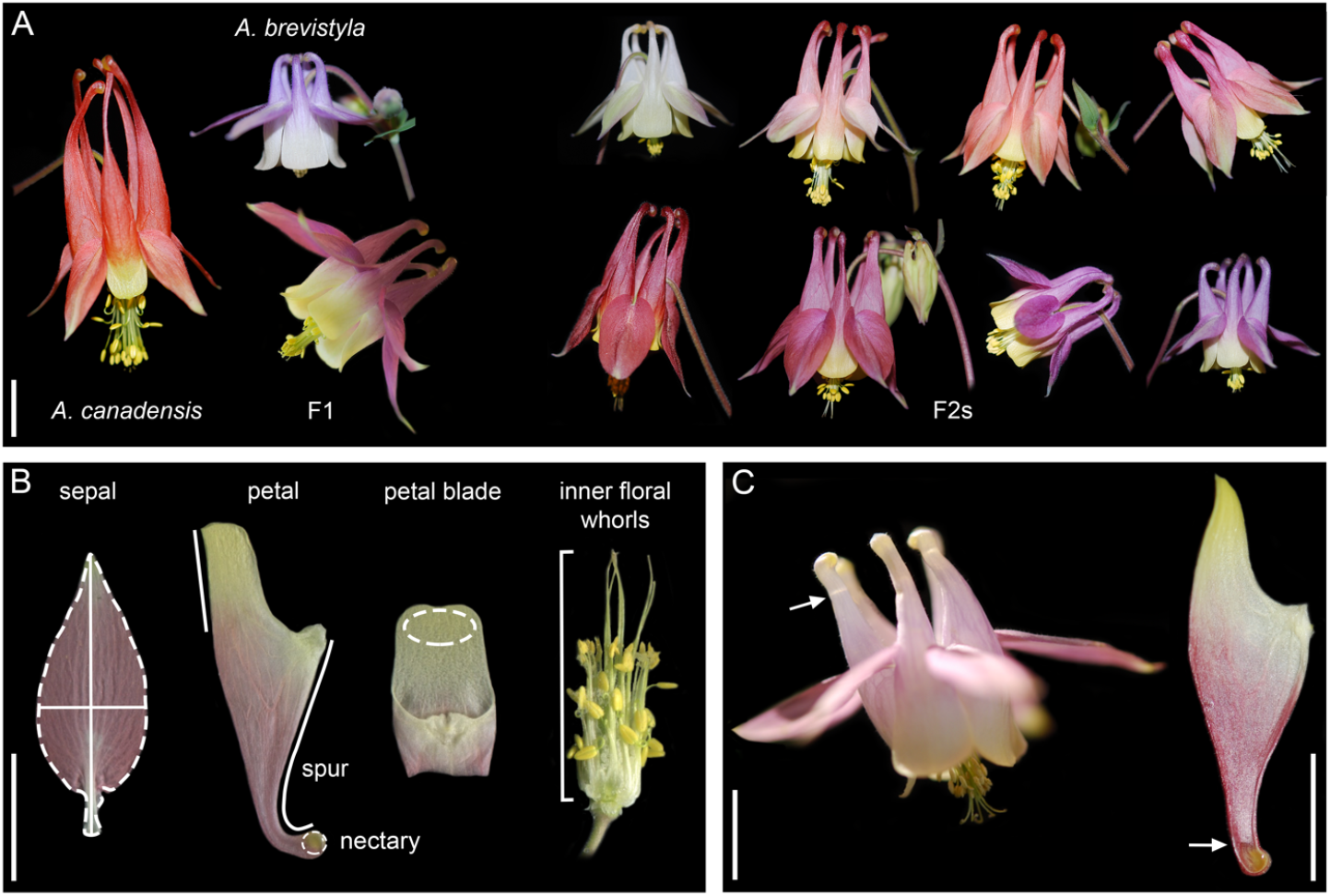
Mapping population and phenotyping. **A.** Parent species *A. canadensis* and *A. brevistyla*, an F1 resulting from their cross, and a selection of the 352 members of the F2 population highlighting its phenotypic diversity. **B.** Morphological and color traits were measured from floral organ scans of three flowers per F2 plant. Example floral organ scans and the phenotypes they produced are shown. Sepal scans were used to determine their area, length, width, and CIE L* a* b* color values; petal scans were used to determine blade length, spur length, spur curvature, and nectary area; petal blade scans were used to determine the blade CIE L* a* b* color values; and scans of the inner whorls of floral organs were used to determine pistil length from attachment point to stigma tip (bracket). **C.** Nectar volume and concentration were measured in all five petals from three flowers per F2 plant, and total sugars were calculated from those values. Petals were dissected longitudinally and nectar (arrows) was collected using a capillary tube. Scale bars = 1cm.

## Materials and Methods

### Genetic cross and growth conditions

*A. brevistyla* seeds were collected from a wild population near Tucker Lake in Alberta, Canada, and *A. canadensis* seeds from a wild population in Ithaca, New York. The parent plants were grown from this seed in the greenhouse facilities at the University of California Santa Barbara. An *A. brevistyla* paternal plant and an *A. canadensis* maternal plant were crossed to produce the F1 generation (Fig. 1A). Five F1 plants were self-pollinated to produce the F2 population. Seeds were shipped to Harvard University, where approximately 2000 seeds were sown in plug trays containing Fafard 3B Soil and stratified at 4° C for 4 weeks. They were then moved to growth chambers programmed to 16 hour days at 18° C, and 13° C nights. 366 plants germinated and were transferred to 6-inch pots and again to gallon pots as they matured. When they had produced 11 leaves, they were vernalized at 4° C for 8 weeks to promote flowering. We staggered stratification, transplanting, and vernalization in order to make phenotyping more manageable. The first batch of plants (143 total, the offspring of two of the five F1s) were grown post-vernalization in the greenhouse facilities at the Arnold Arboretum of Harvard University. The second batch of plants (223 total, the offspring of the other three F1s) were grown post-vernalization in the greenhouse facilities at Harvard University’s main Cambridge campus. Both batches experienced the same light intensity and day length; the Arboretum plants experienced stable temperatures of 18° C during the day and 13° C at night, but the Harvard campus greenhouse experienced warmer temperatures due to limited cooling ability. In total, 352 plants reached the flowering stage.

### Genotyping

A multiplexed shotgun sequencing approach was used to genotype the F2s (Filiault et al. 2018). DNA was extracted from young leaf tissue collected from each F0 parent (flash frozen) and each F2 (desiccated using silica gel) using Qiagen DNEasy reagents and Magattract beads (Qiagen, Inc.). Sequencing libraries for the two F0 parents were prepared using the NEBNext Ultra II kit (NEB) and sequenced to ~40x coverage as 150 bp reads on an Illumina MiSeq at the Biological Nanostructures Lab in the California NanoSystem Institute at UC Santa Barbara. For the F2s, DNA was quantified using a Qubit 2.0 fluorometer (ThermoFisher Scientific), and 100ng of each sample was used to prepare sequencing libraries with the iGenomX RIPTIDE kit. Libraries were pooled and sequenced as 150bp paired-end reads in one lane of an Illumina NovaSeq 6000 by the DNA Technologies & Expression Analysis Core at the UC Davis Genome Center to achieve ~1-2x coverage for each F2.

All sequence reads were aligned to the *A. coerulea* ‘Goldsmith’ v3.1 reference genome (https://phytozome.jgi.doe.gov) using the Burrows-Wheeler aligner (Li and Durbin 2009) as in Filiault et al. (2018). Variable sites in the parents were identified using SAMtools 0.1.19 (Li et al. 2009) and custom scripts were used to identify the positions and genotypes at which the parents were homozygous for different alleles. These sites were used to assign reads in the F2s as having either *A. canadensis* or *A. brevistyla* ancestry. The genome was broken up into windows of either 0.5 Mb or 1Mb, depending on recombination rate as determined in prior crosses, and the frequency of reads with ancestry for each F0 parent was used to determine the genotype of the bin (see Filiault et al. 2018, Ballerini et al. 2020 for details). These bins and genotypes were used as markers to construct a genetic map and conduct QTL mapping (see below).

### Phenotyping

Seventeen traits were phenotyped in the F2s and parents, spanning floral color and nectar composition, as well as sepal, petal, and pistil morphology (Table S1). Phenotypes were collected from nine flowers per plant, divided into three sets of three. The first set was used to phenotype all of the morphological traits, and the flower color found in the sepals and petal nectar spurs. The second set was used to phenotype the petal blade color. The third set was used to phenotype the nectar traits. All flowers were phenotyped at anthesis after approximately half of the anthers had dehisced.

Where possible, the three flowers collected for the first set were the terminal flower and the first two lateral flowers on the primary inflorescence. Sepals were removed and taped to a piece of black paper with their adaxial side up using clear packing tape; petals were removed, folded longitudinally, and also taped to the paper; and the inner floral whorls (stamens, staminodes, pistils) were left intact. These organs were scanned with a ruler using an Epson Perfection V300 scanner at 600 dpi (Fig. 1B). Sepal traits (area, length, width), nectary area, petal blade length, and pistil length were measured from the scans in FIJI (Schindelin et al. 2012). The petal nectar spurs and the sepals are the same color in most North American *Aquilegia* species; therefore, we quantified this color from the sepals only to avoid detecting the contrasting petal blade color, and will hereon refer to it as ‘sepal color.’ To quantify it, the sepals were cropped from the whole scan and the background was removed in Adobe Photoshop; RGB values were measured using MATLAB (2019b) and averaged across all pixels in each scan; and the average RGB values were converted to the three-dimensional CIE L*a*b* color space in R using the convertColor command (R core team 2020). The a* axis represents variation in red-green, b* represents blue-yellow, and L* light-dark.

For nectar spur length and curvature quantification, the xy-coordinates of the petal attachment point and the spur tip were extracted from each petal scan in ImageJ. The petal RGB image was then filtered and binarized using the ‘imgaussfilt’ and ‘imbinarize’ functions in MATLAB. A smooth curve representing the petal spur was then obtained using the ‘bwboundaries’ function in MATLAB, with the extracted feature coordinates used for guiding the segmentation. The spur length was then quantified by the total length of the curve segment between the spur tip and the attachment point. The spur curvature quantification was done by computing the curvature difference between the curve segment of each F2 petal and that of a straight reference petal from an *A. canadensis* flower. For each petal, the signed curvature 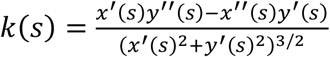 (Do Carmo 2016) was first computed, where the first zero of the curvature near the spur tip was parameterized as *s* = 0 and the attachment point was parameterized as *s* = 1. For the computation of *k*(*s*), the curve segment was rescaled to be with unit length in order to remove the effect of size on the curvature quantification. The spur curvature was then quantified by the *L*^2^-norm of the difference between the signed curvature *k*(*s*) of the F2 petal and that of the straight reference petal, evaluated from *s* = 0 to *s* = 0.5 (ie, the lower half of the spur). Note that the part with *s* ≥ 0.5 (the upper half of the spur) was excluded as the variation in shape near the attachment point is irrelevant to the spur curvature.

The second set of flowers were selected at random from the inflorescence. The petals were taken off the flower and their nectar spurs were removed just below the attachment point. The blades were taped adaxial side up and scanned as for the first batch of flowers. Color was quantified as for the sepals, but instead of sampling across the entire organ, only the distal portion of the blade was used to avoid capturing the spur color, which is identical to the sepal.

The third set of flowers were also selected at random. Each spur was slit longitudinally at the attachment point for easier access to the nectar, which was collected from each petal individually using a capillary tube (Fig. 1C). Height in the tube was measured to the nearest 0.5mm, the nectar was transferred to a hand-held refractometer (Eclipse Low Volume Nectar Refractometer, 0-50 & 45-80, Bellingham & Stanley), and the °Brix measurement (sugar concentration by mass) was recorded. All nectar measurements were taken between 9am and noon to minimize circadian variation in nectar traits. Total nectar volume per flower was calculated from the nectar height measurements and the circumference of the capillary tube, and the sugar concentration data were adjusted based on the temperature of the room at the time the measurement was taken according to the manufacturer’s instructions. While we do not know the sugar type data for *A. brevistyla*, the dominant sugar in *A. canadensis* nectar is sucrose (Macior 1978). Total sugars was calculated in R by fitting a power model to the °Brix and molarity data in the Concentrative Properties of Aqueous Solutions Standard Table for Sucrose (CRC Handbook of Chemistry & Physics 2019); using that model to convert the temperature-adjusted sugar concentration values to molarity; and then multiplying molarity by nectar volume (as in Bolten et al. 1979).

For all traits, the final phenotypic value for each F2 is the average phenotypic value of the three flowers after averaging across multiple organs within a flower, where applicable. Not all 352 F2s were phenotyped for all traits: the first set of flowers were prioritized so that the vast majority of F2s have all morphological traits quantified, whereas only 303 F2s produced enough flowers to quantify nectar traits. Parental phenotypes were also collected as for the F2s; the plants that were phenotyped were not the parents themselves, but close relatives (siblings and offspring). Normality of the phenotypic distributions was assessed with the Shapiro-Wilk Test using the shapiro.test command in R (R core team 2020), and pairwise correlations were tested with the Spearman method using the Hmisc R package (Harrell 2020).

### QTL mapping

All QTL mapping analyses were done using the R package R/qtl v1.46-2 (Broman et al. 2003). The genetic map was estimated from F2 genotypic data, which was then combined with the phenotypic data to perform QTL mapping using the Hayley-Knott regression method. The ‘scanone’ function was used to estimate single QTL locations on each chromosome, and ‘scantwo’ was used to determine interactions between QTL pairs. These data were used as inputs for a multiple QTL model, which was assessed and adjusted using the ‘fitqtl’ and ‘refineqtl’ commands. The percent variation explained (PVE) for each component of the model and for the model as a whole were estimated using these functions as well. 1.5 LOD intervals of each significant QTL were obtained from the refined model. 1,000 permutations were used to calculate a significant LOD cutoff of 3.5, equivalent to a 5% false discovery rate. For the nectar traits (volume, sugar concentration, and total sugars), all analyses were performed with a batch covariate to account for the different growing conditions experienced by the two batches of plants.

### Identification of color candidate loci

Likely homologs for regulatory and biosynthetic genes involved in producing floral pigments were identified by DELTA-BLAST 2.11.0+ (Boratyn et al. 2012) searches of the predicted *Aquilegia* proteome (version 3.1, https://phytozome.jgi.doe.gov), with protein sequences of genes characterized in other taxa as input queries (Table S2). Searches were also run against the predicted *Arabidopsis thaliana* proteome (TAIR10; Berardini et al. 2015). Blast results were then filtered based on percent identity and alignment length.

Gene trees were constructed if more than one candidate *Aquilegia* homolog was identified (Fig. S1). Multiple sequence alignment of characterized genes, *Aquilegia* blast hits, and *Arabidopsis* blast hits was performed with clustal omega, using default parameters (Sievers et al. 2011). Alignments were evaluated in Jalview (Waterhouse et al. 2009). Poorly aligned sequences lacking conserved elements were removed from the alignment, and, except for alignments of MYBs, positions with a quality score less than 50% were masked. MYB alignments were manually trimmed to R2R3 or R3 regions (Stracke et al 2001.) and insertions were masked. Neighbor joining trees were then constructed based on BLOSUM62 distances (Jalview; Fig. S1).

## Results

### Phenotypic data

In our F2 population of 352 individuals, we analyzed measurements of 17 traits that captured floral color, nectar composition, and organ morphology in order to understand trends in phenotypic distribution, relationship to parental phenotypes, and correlations between traits. The three sepal size traits were normally distributed, as were spur length and blade CIE b* and CIE L* (Fig. 2, S2). All other traits were non-normally distributed. In general, the traits exhibited unimodal distributions except for the three sepal color CIE axes and sugar concentration, which were bimodally distributed. We found that roughly half of the F2 trait values fell predominantly between parental means (spur curvature, spur length, blade length, pistil length, sepal and blade CIE b*) while the other half exhibited transgressive segregation (sepal area, length, and width; sepal and blade CIE a* and CIE L*; nectary area; and all of the nectar traits).

**Figure 2.**
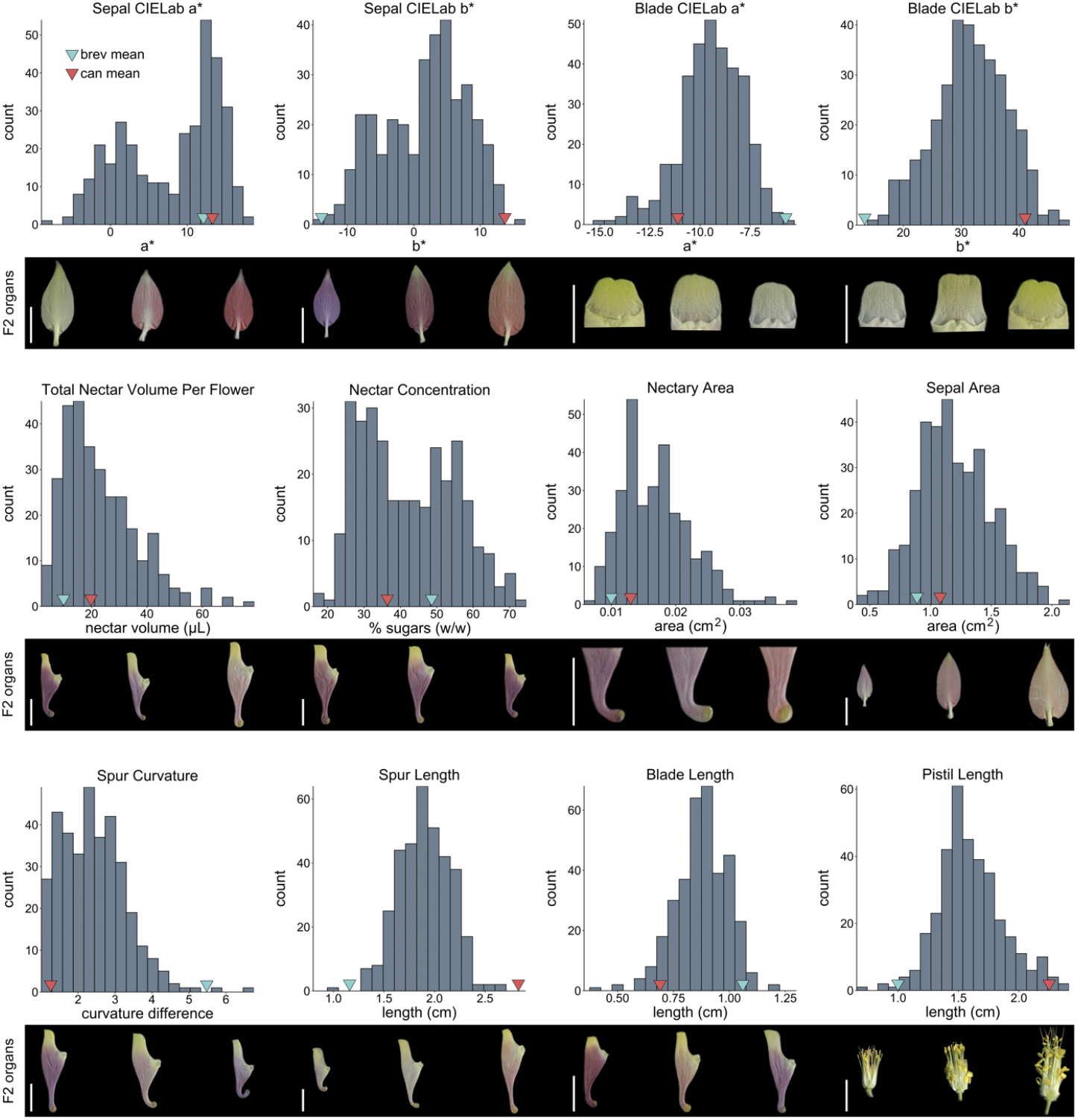
F2 population floral trait histograms (remainder in Fig. S2). Blue and red arrows mark the phenotypic means of *A. brevistyla* (brev mean) and *A. canadensis* (can mean) plants that were closely related to the parents. Representative floral organs from F2s with the lowest, mean, and highest phenotypic values are shown below each histogram. Scale bars = 1cm.

Correlations were common both among and between morphological and biochemical traits in the F2 population, with the largest Spearman correlation coefficients between traits measured within the same organ (Table 1). Sepal color CIE L* and CIE a* topped the list (−0.95, *p* <0.001), followed closely by sepal size traits (sepal area and width, 0.92, *p* <0.001; sepal area and length, 0.87, *p* <0.001). Nectar traits exhibited the next highest correlation coefficients with nectar volume and concentration at −0.80 (*p* < 0.001), and nectar volume and total sugars at 0.79 (*p* < 0.001). The highest correlation in organs of different types was between pistil length and spur length (0.71, *p* <0.001). Although all pairwise combinations of CIE L*a*b* values were significantly correlated for both sepal and blade color, very different correlation patterns emerged in the sepal and blade colors when we plotted these phenotypes. In blades, the phenotypes form a single continuous cluster with yellower blades having low CIE a* and high CIE*b values, and whiter blades having high CIE a* and low CIE b* values (Fig. 3, top). In sepals, however, two clusters are present (Fig. 3, bottom): one in the high CIE b*, low CIE a* quadrant that appear mostly white and yellow, and the other in the high CIE a* half that have pigment ranging from red (high CIE b*) to blue (low CIE b*). Given these patterns, correlations between single CIE axes and morphological traits could be informative for blade color, but not for sepal color, because a high CIE a* value for blades consistently means that the blades are white, while sepals with high CIE a* values could be white, red, or anything in between.

**Figure 3.**
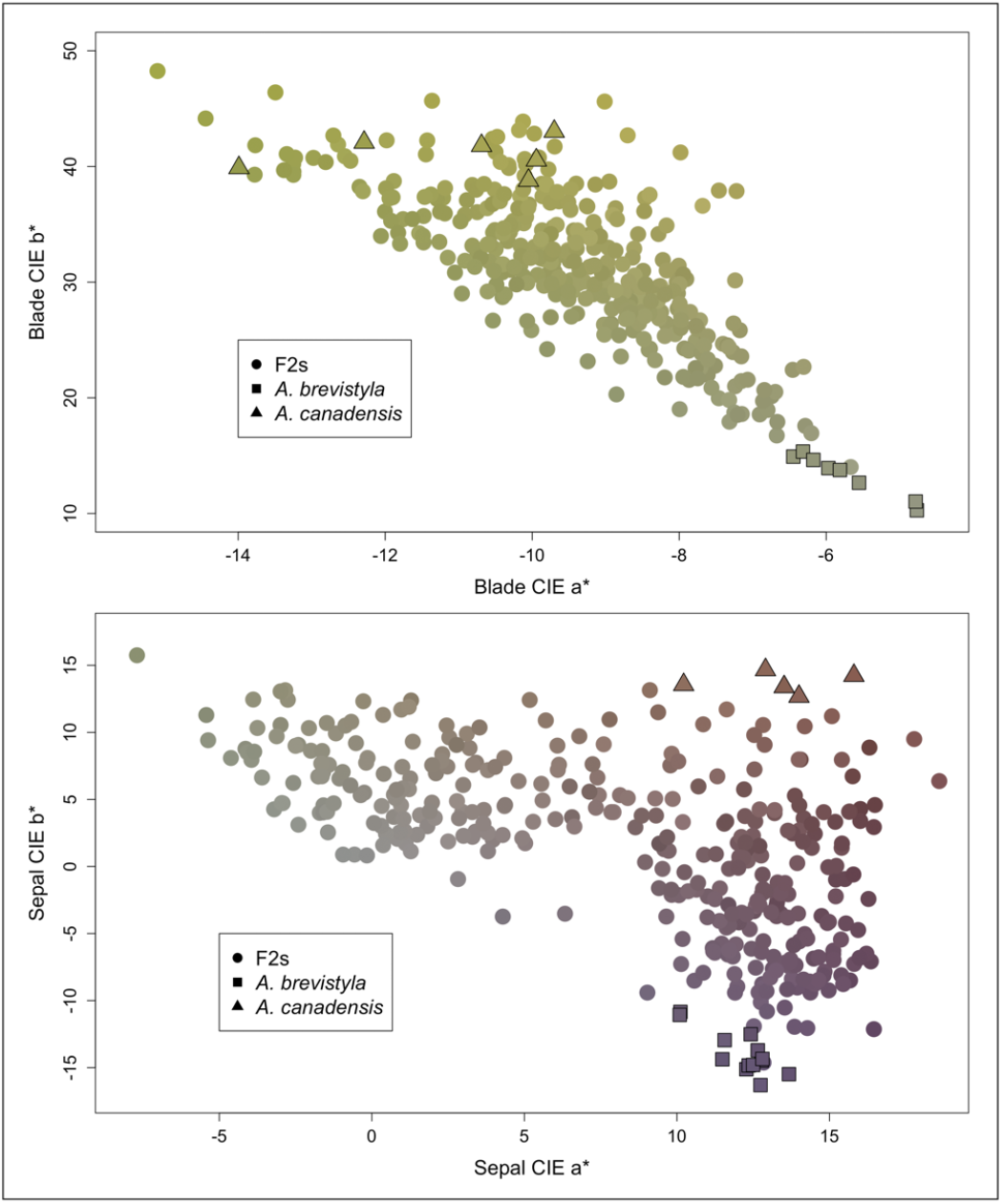
Plots of blade color CIE a* and CIE b* (top) and sepal color CIE a* and CIE b* (bottom), exhibiting different correlation patterns. Points are color-coded based on the mean RGB values of the blade or sepal for that individual F2 or parent plant.

**Table 1.** Pairwise trait correlations. Cells are shaded based on absolute value of ρ (Spearman correlation coefficient). *0.05, **0.01, ***0.001

|  | Sepal CIE L | Sepal CIE a | Sepal CIE b | Blade CIE L | Blade CIE a | Blade CIE b | Nectar vol. | Nectar conc. | Total sugars | Nectary area | Sepal area | Sepal length | Sepal width | Blade length | Spur length | Spur curv. |
| --- | --- | --- | --- | --- | --- | --- | --- | --- | --- | --- | --- | --- | --- | --- | --- | --- |
| Sepal CIE a | -0.95*** |  |  |  |  |  |  |  |  |  |  |  |  |  |  |  |
| Sepal CIE b | 0.60*** | -0.62*** |  |  |  |  |  |  |  |  |  |  |  |  |  |  |
| Blade CIE L | 0.14* | -0.09 | 0.20*** |  |  |  |  |  |  |  |  |  |  |  |  |  |
| Blade CIE a | -0.20*** | 0.17** | -0.40*** | -0.28*** |  |  |  |  |  |  |  |  |  |  |  |  |
| Blade CIE b | 0.19*** | -0.15** | 0.45*** | 0.58*** | -0.73*** |  |  |  |  |  |  |  |  |  |  |  |
| Nectar vol. | 0.02 | -0.03 | 0.10 | 0.05 | -0.15** | 0.12* |  |  |  |  |  |  |  |  |  |  |
| Nectar conc. | -0.02 | 0.05 | -0.08 | 0.02 | 0.06 | 0.05 | -0.8*** |  |  |  |  |  |  |  |  |  |
| Total sugars | 0.07 | -0.04 | 0.10 | 0.13* | -0.22*** | 0.29*** | 0.79*** | -0.31*** |  |  |  |  |  |  |  |  |
| Nectary area | 0.03 | -0.06 | 0.25*** | 0.22*** | -0.36*** | 0.31*** | 0.62*** | -0.48*** | 0.49*** |  |  |  |  |  |  |  |
| Sepal area | -0.16** | 0.16** | 0.16** | 0.10 | -0.32*** | 0.26*** | 0.58*** | -0.45*** | 0.47*** | 0.73*** |  |  |  |  |  |  |
| Sepal length | -0.17** | 0.16** | 0.03 | -0.04 | -0.15** | 0.03 | 0.55*** | -0.44*** | 0.42*** | 0.64*** | 0.87*** |  |  |  |  |  |
| Sepal width | -0.25*** | 0.25*** | 0.18** | 0.18*** | -0.39*** | 0.39*** | 0.45*** | -0.33*** | 0.38*** | 0.65*** | 0.92*** | 0.67*** |  |  |  |  |
| Blade length | -0.21*** | 0.18*** | -0.38*** | -0.40*** | 0.43*** | -0.55*** | 0.02 | 0.02 | 0.00 | 0.04 | 0.09 | 0.34*** | -0.09 |  |  |  |
| Spur length | 0.05 | -0.01 | 0.40*** | 0.40*** | -0.47*** | 0.59*** | 0.24*** | -0.19*** | 0.22*** | 0.46*** | 0.50*** | 0.23*** | 0.60*** | -0.45*** |  |  |
| Spur curv. | -0.04 | 0.06 | -0.14** | -0.07 | 0.13* | -0.08 | -0.20*** | 0.21*** | -0.11 | -0.34 | -0.21*** | -0.15** | -0.20*** | 0.18*** | -0.37*** |  |
| Pistil length | 0.02 | 0.00 | 0.42*** | 0.21*** | -0.32*** | 0.35*** | 0.31*** | -0.23*** | 0.25*** | 0.60*** | 0.54*** | 0.36*** | 0.57*** | -0.24*** | 0.71*** | -0.5*** |

### Genotypic data

Sequence reads at 200,619 SNP positions across 671 marker bins, averaging 299 SNP positions per bin, were used to genotype the F2 population. Several bins had very few SNP positions (< 33 loci/bin) making genotyping difficult, with the result that only 620 markers were used to estimate the genetic map. Genetic ordering of markers is largely consistent with the physical assembly of the *A. coerulea* “Goldsmith” reference genome sequence, however, similar to what has been seen in other genetic crosses in *Aquilegia*, several markers mapped to genetic locations inconsistent with the physical map, suggesting that there may be some physical differences between the individual used for assembling the reference genome and other species (Fig. S3; Filiault et al., 2018, Ballerini et al., 2020). Overall, patterns of recombination across chromosomes in this cross mimic those seen in other *Aquilegia* crosses, whereby recombination rate is much higher at chromosome tips but rarely occurs across large physical spans in the middle of chromosomes (Fig. S3). The cross exhibits a substantial amount of transmission ratio distortion, overall favoring *A. canadensis* alleles, possibly because *A. canadensis* pollen has evolved to travel down longer styles than that of *A. brevistyla*. Loci homozygous for *A. canadensis* alleles are greatly overrepresented across most of chromosome 1 (chr1), and moderately overrepresented in regions of chrs 3, 5, and 6 (Fig. S4). *A. brevistyla* homozygotes are strongly overrepresented on one end of chr2 (Fig. S4).

### QTL mapping

We identified a total of 66 QTL for the 17 traits mapped in this study, with 2-6 significant QTL per trait (Table 2, Fig. 4, S5). However, not all 66 QTL are unique; for example, all three CIE axes for sepal color have the same significant QTL at 48.3cM on chr1. We define a major effect QTL as one that explains 25% or more of the variance in the F2 population (Bradshaw et al. 1995), and a QTL of moderate effect as one with 10-25% variance explained (PVE). With this framework in mind, a single major locus and four minor loci control the sepal color (CIE L*a*b* as a whole), while QTL on every chromosome contribute to blade color and vary from minor to moderate, and moderate to major, depending on which axis of the CIE L*a*b* color space is being mapped. The multiple QTL model for sepal color had the highest overall PVE in the study (87 for CIE L*, 81.7 for CIE a*, 87.3 for CIE b*). The QTL for the nectar traits had the lowest overall PVEs, with each explaining less than 10% of the variance in the F2s. We included a batch covariate in the QTL models for nectar traits to account for the different growth conditions in the two greenhouses; this covariate explained 35% of the variance in nectar volume and 51.2% of the variance in concentration, but only 5.8% of the variance in total sugars. The multiple QTL models for morphological traits had overall PVEs ranging from 22.6 for sepal length, to 74.4 for pistil length, and had equally variable genetic architectures: a single major locus and two minor ones contribute to spur curvature, while the maps of blade length and spur length contain six loci of minor to moderate effect. No QTL interactions were detected except in the maps for sepal color, where the major locus on chr1 interacts with the minor loci on chrs 2, 4, and 6.

**Figure 4.**
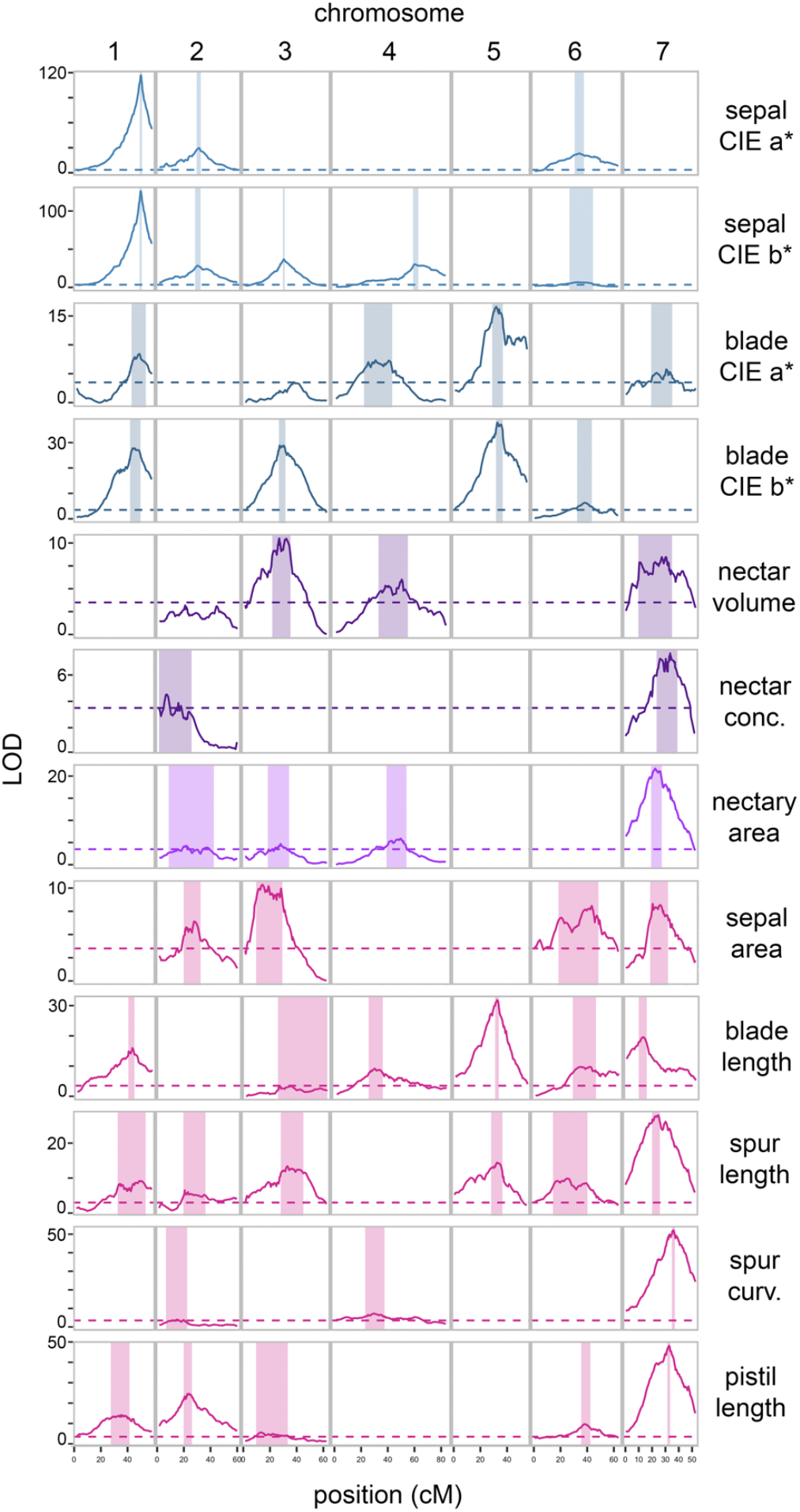
Floral trait QTL maps (remainder in Fig. S5). Dashed line represents the significant LOD cutoff of 3.5, shaded areas represent the 1.5 LOD interval for each peak. Abbreviations: conc., concentration; curv., curvature.

**Table 2.**
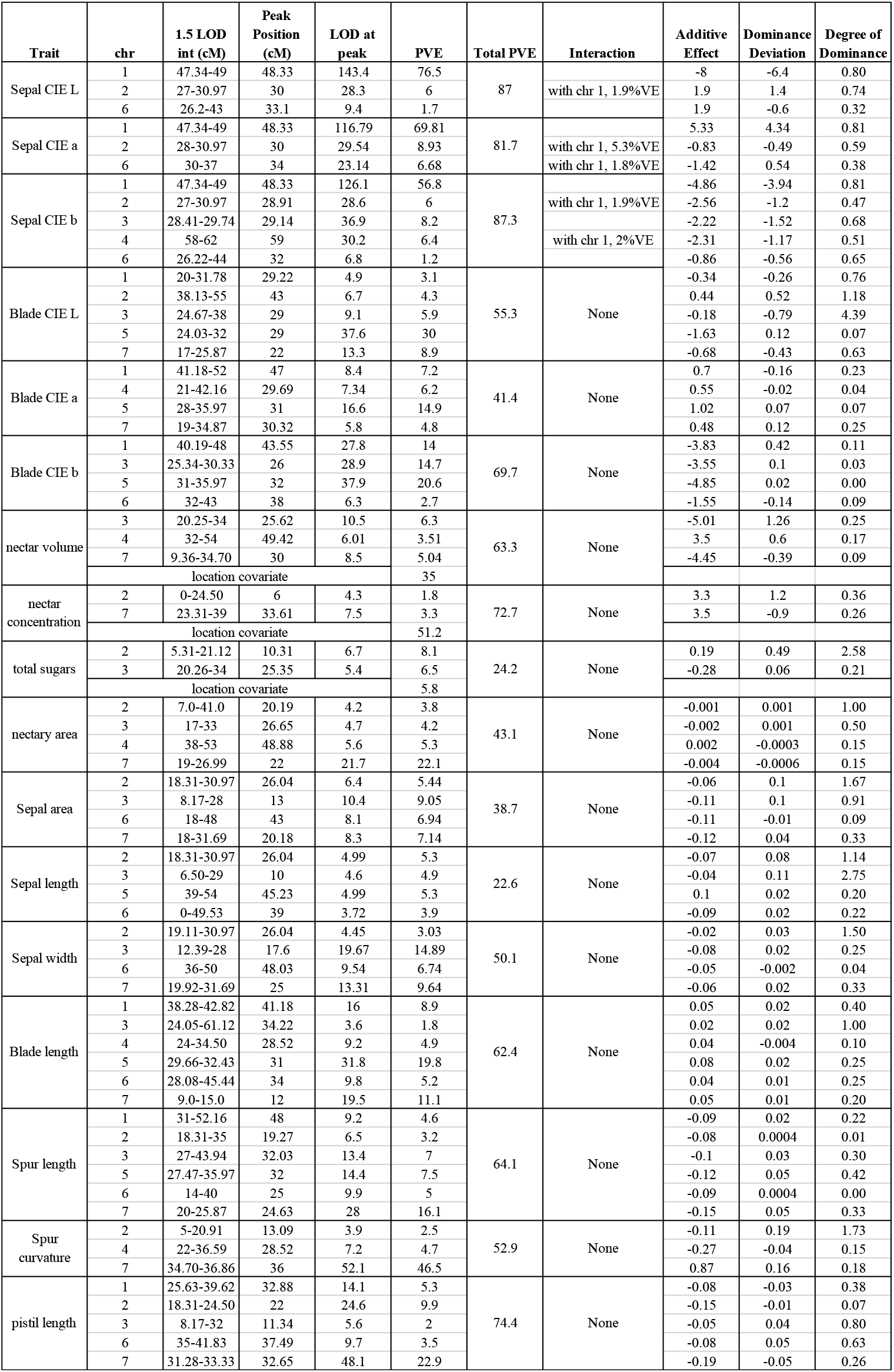
QTL data for all traits. Abbreviations: chr, chromosome; int, interval; PVE, Percent Variance Explained.

All alleles at identified QTL were in the direction of parental divergence except for single loci in the maps for sepal length (chr5), sepal CIE a* and CIE L* (chr1), blade CIE L* (chr2), spur curvature (chr4), nectar volume (chr4), total sugars (chr2), and nectary area (chr4). All transgressive traits except sepal color exhibited overdominance at one or more loci, most notably the locus on chr2 for total sugars and chr3 for sepal length.

QTL co-localization, defined by overlapping 1.5 LOD intervals, was abundant in this dataset (Fig. 4). All of the sepal size loci co-localize, with the exception of the unique chr5 locus for sepal length. The loci on chrs 2, 6, and 7 for sepal size overlap with sepal color loci. Of the six QTL identified for blade length, four overlap with loci for blade color and spur length. The three nectar composition traits and nectary area each have 2-3 peaks on chrs 2, 3, 4, or 7, all of which co-localize. Notably the large peaks on chr7 for spur length and spur curvature are not colocalized, although their peaks are less than 12cM apart. The QTL on chrs 1, 2, 3, and 6 are colocalized between spur length and pistil length, and their large peaks on chr7 are separated by 5cM. The nectar peaks co-localize with those of spur length and pistil length on chrs 2, 3, and 7.

### Identification of potential causative genes under QTL peaks

While many genes related to organ growth and development are beneath the 1.5 LOD intervals of the morphological trait QTL, we will not speculate about potential causative loci here because characterization of the developmental basis of these traits is still ongoing and not all of these genes have been functionally validated in other systems. The genetic basis of floral color, on the other hand, has been well-characterized in many systems (reviewed in Tanaka et al. 2008), and based on this understanding we built an extensive catalog of likely homologs of floral pigment-related genes (Table S3-S4, Fig. S1). The molecular underpinnings of nectar synthesis are becoming clearer, with multiple key genes having been recently characterized (Chen et al. 2010; Lin et al. 2014; Min et al. 2019). We identified many of these known genetic players in color biosynthetic pathways and nectar composition in the 1.5 LOD intervals of QTL contributing to these respective traits.

Floral pigment analyses done by Taylor (1984) identified carotenoids and anthocyanins in *A. canadensis*, and anthocyanins (but no carotenoids) in lavender-blue species similar to *A. brevistyla*. We identified *Aquilegia* homologs of known members and regulators of the anthocyanin biosynthetic pathway (ABP) and the carotenoid biosynthetic pathway (CBP) beneath peaks in the sepal color maps (sepal CIE L*, a*, and b* maps collectively). The blade color maps only contained CBP *Aquilegia* homologs, consistent with the lack of anthocyanins in the blades of the parental species.

Two core members of the ABP and a family of modification enzymes appear to be controlling anthocyanin production in this cross (Tanaka et al. 2008; Matsuba et al. 2010; Table S3). *FLAVONOID 3 ′5 ′-HYDROXYLASE (AqF3′5 ′H)*, a major branching enzyme in the ABP, is under the large-effect QTL on chr1; several duplicates of *GLYCOSYL HYDROLASE (AqGH)*, a vacuolar anthocyanin modification enzyme, are under the chr2 locus; and *DIHYDROFLAVONOL REDUCTASE (AqDFR)*, another member of the ABP downstream of *F3¢5¢H*, is under the peak on chr6. The minor peak on chr4 is the only sepal color QTL for which we could not identify a gene known to be involved in the ABP.

Thirty genes – core members of the CBP, or peripheral players and regulators thereof – are found in the 1.5 LOD intervals of the color QTL and could be candidates for contributing to carotenoid dynamics in this cross. CBP regulators *CAROTENOID CLEAVAGE DIOXYGENASE 4.1 and 4.2 (AqCCD4.1* and *Aq CCD4.2)* are under the chr3 locus shared between the sepal and blade color maps (Ohmiya 2009), as is *ORANGE PROTEIN (AqOR)*, a gene that affects carotenoid accumulation (Li et al. 2001). Except for the minor locus on chr4, all of the QTL specific to blade color each contain 1-5 genes known to be involved in carotenoid synthesis or regulation (summarized in Table S4).

The chr2 locus that is shared between nectar concentration, total sugars, and nectary area contains *AqLRP*, a member of the *STYLISH (STY)* gene family that is necessary for nectary development in the Ranunculaceae (Min et al. 2019). The locus on chr3 in the nectar volume, total sugars, and nectary area maps contains an *Aquilegia* homolog of a member of the *SWEET* family, bidirectional sugar transporters involved in nectar secretion (Chen et al. 2010). The chr7 and chr4 loci each contain several sugar transporters and sucrose synthesis genes; unique to the sugar concentration chr7 1.5 LOD interval is *SUCROSE PHOSPHATE SYNTHASE-1 (SPS1F)*, a sucrose synthesis gene that is necessary for nectar production in *Arabidopsis* (Lin et al. 2014).

## Discussion

We identified QTL for multiple complex traits related to pollination syndromes in *A. canadensis* and *A. brevistyla*. These traits cut across well-defined genetic pathways such as anthocyanin biosynthesis, as well as those that are less understood such as organ size and shape.

### Color

#### Anthocyanin pathway

*A. canadensis* flowers make red cyanidin and pelargonidin anthocyanidin pigments, while lavender-blue species like *A. brevistyla* only make delphinidins (Taylor 1984). The QTL maps for sepal color consist of a single locus of major effect that interacts with several loci of minor effect: we have identified *AqF3′5′H* at the large QTL on chr1, and *AqGH* and *AqDFR* at the minor loci on chrs 2 and 6, respectively, as good candidate loci for controlling color variation. Here we present a model for how the interaction of these candidate loci may contribute to sepal pigmentation in this cross, which categorizes the F2s into three classes based on their phenotypes and genotypes at these loci (summarized in Fig. 5).

**Figure 5.**
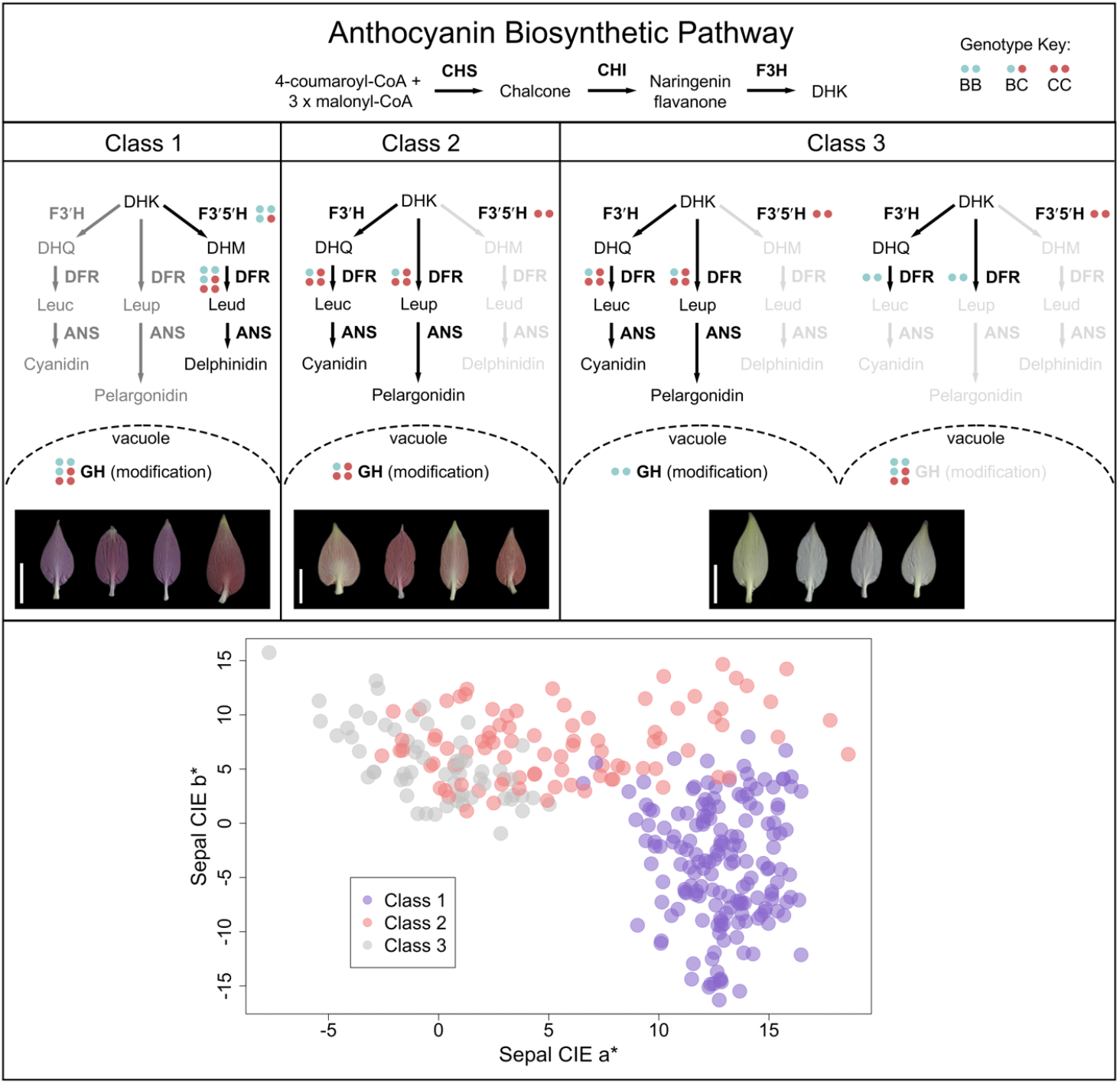
Model of anthocyanin dynamics in this cross depicting three classes of pigmentation in the F2s. The header panel shows the first steps of the core ABP, while each Class panel (1-3) shows the later branching steps of the ABP and downstream GH modification, which occurs in the vacuole. Enzymes are in bold and ABP branches that experienced reduced or no flux are in dark grey and light grey, respectively. Allelic genotypes of key enzymes are represented by pairs of blue (*A. brevistyla* allele) and red (*A. canadensis* allele) dots. The left Class panel shows Class 1 of F2 anthocyanin pigmentation; F2s with an *A. brevistyla F3′5′H* allele are all pigmented blue-purple due to flux primarily moving through the delphinidin branch of the pathway, and all *AqDFR* and *AqGH* genotypes accepting substrates from that branch. The middle Class panel shows Class 2: F2s homozygous for the *A. canadensis F3′5′H* allele are pigmented pink-red because flux is moving through the pelargonidin and cyanidin branches, and they have *A. canadensis AqDFR* and *AqGH* alleles that are compatible with those substrates. The right Class panel shows Class 3: white and yellow F2s lack anthocyanins because they are homozygous *A. canadensis* at *AqF3′5′H* and homozygous *A. brevistyla* at either *AqDFR* or *AqGH*, which can only process substrates from the delphinidin branch. Representative sepals from each class of pigmentation and corresponding genotypes are shown below the ABP summaries in each Class panel. Scale bar = 1cm. The bottom panel shows how these three classes cluster in the CIE a* and b* color space. CHS, chalcone synthase; CHI, chalcone isomerase; F3H, flavanone 3-hydroxylase; DHK, dihydrokaempferol; F3′H, flavonoid 3’-hydroxylase; F3′5′H, flavonoid 3’,5’-hydroxylase; DHQ, dihydroquercitin; DHM, dihydromyricetin; Leuc, leucocyanidin; Leup, leucopelargonidin; Leud, leucidelphinidin; DFR, dihydroflavonol reductase; ANS, anthocyanidin synthase; GH, glycosyl hydrolase.

*AqF3′5′H* is likely the primary cause of the shift from blue to red in this cross, and has also been implicated in blue-to-red color shifts in *Phlox, Penstemon*, and *Iochroma* (Hopkins and Rausher 2011; Smith and Rausher 2011; Wessinger and Rausher 2013). Expression of *F3′5′H* diverts anthocyanin precursors down the branch of the ABP that produces blue delphinidins, while lack of *F3′5′H* expression yields red pelargonidins and cyanidins (reviewed in Tanaka et al. 2008). Class 1 F2s are defined by having at least one *A. brevistyla AqF3′5′H* allele, and clustering in the high CIE a*, low CIE b* range of the sepal color space (Fig. 5, Class 1 and lower panel). Class 1 F2s are all blue-purple in color, and we believe they are primarily producing delphinidin pigments due to *F3′5 ′H’s* ability to outcompete *F3′H* for substrate (Gerats et al. 1982; Fig. 5, Class 1). Because *A. canadensis* does not make delphinidins, we assume that its *F3′5′H* allele is either non-functional or not expressed, and that in plants homozygous for *A. canadensis* at *AqF3′5′H*, flux is directed down the other branches of the ABP.

However, there is substantial phenotypic variation among F2s homozygous for *A. canadensis* at *AqF3′5′H*: some are pigmented pink-red and have varying CIE a* values but cluster in the high CIE b* range, while others appear to lack anthocyanins and are pure white or yellow, with low CIE a* and high CIE* b values (Fig. 5, bottom panel). We divide these F2s into Class 2 (pigmented pink-red) and Class 3 (white or yellow), and propose that *AqF3’5H* epistatically interacts with *AqDFR* and *AqGH* to produce these transgressive Class 3 phenotypes. All Class 2 F2s have at least one *A. canadensis* allele at both *AqDFR* and *AqGH*, while Class 3 F2s are homozygous *A. canadensis* at either or both loci (Fig. 5, Class 2 and Class 3).

*DFR* acts downstream of the flavanone hydroxylases (*F3H, F3′H*, and *F3′5′H*) to reduce dihydroflavonols into leucoanthocyanidins, and can have strict substrate specificity for a particular dihydroflavanol (i.e., dihydrokaempferol (DHK), dihydroquercitin (DHQ), or dihydromyricetin (DHM); reviewed in Tanaka et al. 2008). All F2s that are homozygous for *A. canadensis* alleles at *AqF3′5′H* and homozygous for *A. brevistyla* alleles at *AqDFR* appear to lack anthocyanins (Fig. 5, Class 3; Fig. S6), suggesting that the *A. brevistyla* protein is unable to process DHQ or DHK, or is outcompeted by other flavonoid pathway enzymes, as has been shown in *Petunia* and*Mimulus* (Davies et al. 2003; Yuan et al. 2016). In addition to color variation produced by generating different anthocyanidin pigment types, further diversity of floral color is in large part due to modification of anthocyanidin pigments with glycosyl, acyl, and methyl groups. The *AqGHs* on chr2 are homologues of one such modification enzyme, known to act in the vacuole to glucosylate anthocyanidins in *Carnation* as well as *Delphinium*, which is in the same family as *Aquilegia* (Matsuba et al. 2010; Table S3). In this cross, *A. canadensis AqGH* alleles result in redder sepals than *A. brevistyla* alleles, regardless of genotype at *AqF3′5′H* and resulting anthocyanin type (Fig. S7). However, flowers that are homozygous *A. canadensis* at *AqF3′5′H* and have at least one *A. canadensis* allele at *AqDFR*, but are homozygous *A. brevistyla* at *AqGH*, are white-yellow. (Fig. 5, Class 3; Fig. S8). In this case, anthocyanidins were presumably formed and transported into the vacuole, but the *A. brevistyla AqGH* proteins somehow fail to prevent pigment degradation. The GH family is involved in pigment stability (Sasaki and Nakayama 2015), but we do not have enough information to speculate about an exact mechanism in this cross. Broadly, we can say that in our model, *A. brevistyla AqDFR* and *AqGH* alleles are only able to act on dihydroflavonols and anthocyanidins produced by the *AqF3′5′H* branch, whereas *A. canadensis* alleles have less specificity and are able to act on all precursors and pigments regardless of which ABP branch produced them.

The discreet, stepwise nature of the ABP means that a change in a single gene can result in a dramatic change in phenotype, as in *Penstemon*, where *F3′5′H* alone is responsible for the blue-to-red transition in the genus (Wessinger et al. 2014). The genetic architecture of blue-to-red shifts can be more complicated, however, as in *Iris*, where 8 loci contribute to floral color (Brothers et al. 2013). The color shift in the present study is similar to that of *Iochroma*, which also involves *F3′5′H* and a change in substrate specificity at *DFR* (Smith and Rausher 2011), but in *Iochroma* it was not certain if the change in *DFR* was absolutely necessary for the transition from blue to red. In this cross, *AqDFR* and the *AqGHs* did need to lose their substrate specificity during the transition to hummingbird pollination in order for *A. canadensis* to evolve its red pigmentation.

#### Carotenoid pathway

*A. canadensis* flowers contain yellow-orange carotenoid pigments, and species with white petal blades like those of *A. brevistyla* do not (Taylor 1984). While carotenoids are very apparent in the bright yellow petal blades of *A. canadensis*, they also contribute to the red coloration throughout the rest of the flower; indeed, CBP genes appear in the QTL maps for sepal color. The combination of anthocyanins and carotenoids underlying red floral color has been found in other systems including *Mimulus* and Solanaceae (Bradshaw and Schemske 2003; Ng and Smith 2016).

Carotenoid genetic architecture in this system is more complicated than that of the anthocyanins, consisting of multiple loci of small to moderate effect, many of which themselves contain multiple genes known to be involved in the CBP (Table S4). The pleiotropic nature of the CBP and the gaps in our knowledge about carotenoid regulation make it difficult to synthesize these loci into a detailed model such as that presented for the ABP above (Tanaka et al. 2008; Zhu et al. 2010). The sheer number of CBP candidate genes we identified in the 1.5 LOD intervals of blade color QTL precludes us from mentioning them all, so we will focus our discussion on the largest peaks on chrs 5, 1, and 3.

The largest-effect peak is on chr5 and contains *AqZEP*, which acts late in the CBP to convert zeaxanthin into other yellow and orange pigments (Zhu et al. 2010). Silencing of *ZEP* in California poppy results in zeaxanthin accumulation in the petals far exceeding that in wild type plants and an accompanying shift in color from bright orange to yellow (Zhou et al. 2018). However, *AqZEP* is at the edge of the 1.5 LOD interval for the chr5 QTL; closer to the peak marker bin are CBP precursors *HYDROXYMETHYLBUTENYL DIPHOSPHATE REDUCTASE (AqHDR)* and several copies of *GERANYLGERANYL DIPHOSPHATE SYNTHASE* (*AqGGPPS*; Table S4). In addition to carotenoid pigments, these genes generate precursors for metabolites essential for plant functioning such as chlorophyll and various hormones (Beck et al. 2013; Botella-Pavía et al. 2004). Due to the high pleiotropic potential of single-copy *AqHDR* we think it an unlikely candidate for CBP regulation in this cross, but given the multiple copies of *AqGGPPS*, it is possible that one is functioning specifically in floral tissue (Ament et al. 2006).

The CIE a* and CIE b* blade color maps have a peak on chr1 that contains a *WRKY* that has been found to positively regulate *CCD4* in *Osmanthus* (Han et al. 2016); a potential shared carotenoid regulatory mechanism between such disparate angiosperm genera is fascinating and will require more investigation*. AqCCD4.1* and *AqCCD4.2*, as it happens, are underneath the peak on chr3 that is shared between the blade and sepal color maps. *CCD4* regulates carotenoid accumulation by cleaving the pigments into apocarotenoids (Ohmiya 2009), and has been implicated in white/yellow petal color changes in multiple systems, primarily from horticultural cultivars and food crops such as *Petunia* (Kishimoto et al. 2018), azalea (Ureshino et al. 2016), and Chinese kale (Zhang et al. 2019). Also in the chr3 1.5 LOD interval and closer to the peak marker bin than *AqCCD4.1/2* is the *Aquilegia* homolog of *OR*, which was first characterized in cauliflower: a mutation in the gene causes the curd and other vegetative tissues to accumulate large amounts of ß-carotene (Li et al. 2001). The exact mechanism has not been determined but *OR* is thought to be involved in the formation of a lipoprotein structural sink for the pigments.

### Nectar

Studies into the genetic basis of nectar and nectaries are becoming more common, but the fact that floral nectaries have evolved multiple times across the angiosperms makes it difficult to assess the extent to which their genetic underpinnings are conserved (reviewed in Roy et al. 2017). Another complicating factor is that the quantity and composition of floral nectar is determined by the interaction of development, physiology, and the environment. If we start by considering the parents, *A. brevistyla* and *A. canadensis*, the pollination syndrome literature would lead us to expect that the bee-pollinated species would produce lower-volume, higher-concentration nectar compared to the hummingbird-pollinated sister species (Baker 1975; Gegear et al. 2017). While our parental phenotype data seem to align with this trend, we hesitate to say that it is a definitive example due to the fact that the parents experienced different growing conditions, and nectar is sensitive to environmental variation (Canto et al. 2007). All of the *A. canadensis* plants were grown in the cooler-temperature batch, while the *A. brevistyla* plants were split between the two batches and experienced both high and low temperatures. This is the first study to characterize *A. brevistyla* nectar, but the data from the existing *A. canadensis* nectar literature indicate that nectar volume and sugar concentration can vary widely within and between populations (Mavraganis 1998; Noutsos et al. 2015). Sampling, quantification, and analysis methods are also inconsistent between these studies, indicating a need for standardization of nectar phenotyping practices if we want to be able to make meaningful comparisons across species and studies in the future.

In the F2s, the transgressive nature of all four nectar traits suggests that their reshuffled parental alleles released constraints on nectar production present in *A. canadensis* and *A. brevistyla*. While there is not a clear pattern of allelic combinations across QTL that correspond to the highest and lowest trait values, the F2s towards the phenotypic maxima and minima do have novel combinations of parental alleles. The environmental variation experienced by the F2s in different growing conditions had a large effect on the nectar traits: the batch covariate of the QTL model explained 35% of the variance in nectar volume, 51.2% of the variance in sugar concentration, and 5.8% of the variance in total sugars. Despite this, we still found significant QTL for each trait, suggesting that they are genetically robust even under extreme environmental variation. The overall PVE of the QTL model for nectary area (43.1) falls in the middle of the other morphological traits in the study, and has one peak of moderate effect. All nectar traits are significantly correlated with each other in the F2 population, so it comes as no surprise that all nectar trait QTL are co-localized with at least one other’s QTL. However, not all QTL are shared among the four traits, so we must examine each trait’s relationship to the others in the context of their shared and unique loci.

Nectar volume and nectary area have a correlation coefficient of 0.62 (*p* <0.001), and their QTL on chrs 3, 4, and 7 have overlapping 1.5 LOD intervals and peaks within a few cM of each other. Nectary area has a QTL on chr2 that appears unique, but the single QTL analysis of nectar volume detected a QTL on that chromosome as well, which only just missed the significant LOD cutoff in the multiple QTL model (Fig. 4, nectar volume). Therefore, we believe that nectar volume in this system is in large part controlled by the amount of secretory tissue present in a nectar spur. Nectary area and nectar volume are strongly correlated in an F2 population of a cross between bee-pollinated *Penstemon amphorellae* and hummingbird-pollinated *P. kunthii*, and nectary area is predictive of pollination syndrome throughout the genus (Katzer et al. 2019). QTL mapping of nectar traits in *Ipomopsis* also found significant correlation between nectar volume and corolla tube size as well as co-localization of their QTL (Nakazato et al. 2013). These authors hypothesized that this could be due to the effect of nectary area, although they did not measure nectary area directly. It would be fascinating to know if this pattern holds true in other systems and represents a common mechanism for increasing nectar volume during pollinator transitions. The morphology of the *Aquilegia* nectar spur lends itself to relatively easy quantification of nectary tissue area, but phenotyping in other species can be quite complex – *Penstemon* required dissection and microscopy, for example (Katzer et al. 2019). Nonetheless, we encourage this extra effort in future studies, for it could greatly increase our understanding of nectar biosynthesis and evolution in the angiosperms.

Nectar volume and sugar concentration have a correlation coefficient of −0.80 (*p* <0.001), but their relationship is more complicated than that of nectar volume and nectary area. On chr7, there is a peak for nectar volume and sugar concentration, but not total sugars. Homozygous *A. canadensis* F2s at that locus have a higher nectar volume than their homozygous *A. brevistyla* counterparts, but also a lower sugar concentration, indicating that the increase in volume is having a diluting effect on the nectar solutes. A very different scenario is playing out on chr3, however, where there is a peak for nectar volume and total sugars, but not for sugar concentration. Homozygous *A. canadensis* F2s at that peak have both a higher nectar volume and total sugars than homozygous *A. brevistyla* F2s, suggesting that whatever mechanism is controlling volume at this locus is not having a diluting effect on the sugar.

There are multiple sugar transport and synthesis genes under each of these peaks, but we do not have the power to distinguish between a single causative locus or multiple tightly-linked ones, and substantial investigation would be required to determine if these candidates are involved in *Aquilegia* nectar production. Overall, the genetic architecture of nectar traits in *Aquilegia* is similar to that of *Ipomopsis*, in which several loci of small effect contribute to nectar volume (Nakazato et al. 2013). This is in stark contrast to *Penstemon, Petunia*, and *Mimulus*, where QTL mapping of nectar traits identified single large-effect loci critical to pollinator shifts (Bradshaw et al. 1995; Stuurman et al. 2004; Wessinger et al. 2014). We do know from the *Penstemon* study that the loci controlling nectar volume and concentration can be totally separate (Wessinger et al. 2014), while in *Mimulus* they overlap entirely (Bradshaw et al. 1995); *Aquilegia* appears to be somewhere in the middle.

### Morphological traits

Two phylogenies support the sister relationship of *A. brevistyla* and *A. canadensis*, with divergence estimated to have occurred less than three million years ago (Whittall and Hodges 2007; Fior et al. 2013). Major morphological shifts were required during this transition from bee to hummingbird pollination in order to facilitate successful pollen transfer and pollinator reward. At first glance, *A. canadensis* flowers are much larger than those of *A. brevistyla*, but closer examination of each floral whorl reveals a more complicated picture. The sepals, which are petaloid in *Aquilegia* and contribute to pollinator attraction, are actually of a comparable size between the two parents (Fig. 2). The extreme transgressive segregation of sepal size traits in the F2s suggests that the mechanisms maintaining the parents’ modest sepal size are not the same and are released in the F2 generation when recombined. The dramatic morphological differences between the parental flowers are concentrated in the inner whorls. *A. canadensis* petal spurs are on average more than twice as long as *A. brevistyla* spurs and are of a very different shape, but their blades are only half as long. We know from previous work that these two regions of the petal are highly differentiated in terms of gene expression patterns (Yant et al. 2015), and in this study we found variable genetic architectures among the three petal traits. Even though the parental spur sizes are very different, their nectary area is similar; nectary area does however segregate transgressively in this cross, in contrast to the other three petal traits that segregate within their parental extremes. Pistils are also more than twice as long in *A. canadensis* than *A. brevistyla*, likely an adaptation to promote pollen transfer from the head of a hovering hummingbird. A fundamental question in this system (and in the radiation of *Aquilegia* writ large) is how these extreme morphological changes evolved so rapidly.

The abundant co-localization of morphological trait QTL (along with color and nectar traits) could hold part of the answer. It certainly explains large correlation coefficients between many phenotype pairs in the F2 population. Genetic correlation, either due to pleiotropy or tight genetic linkage, can facilitate adaptation and speciation because selection on one trait can easily lead to a correlated response in the other trait (Via and Hawthorne 2005; Hermann and Kuhlemeier 2011). Four of the five peaks in the pistil length map co-localize with those in the spur length map, with the exception of the large QTL on chr7, which are only 8cM apart. Four of the blade length loci co-localize with these traits as well, and all three traits are in the direction of divergence: F2s that are homozygous *A. canadensis* at these loci have longer spurs, longer pistils, and shorter blades than their homozygous *A. brevistyla* counterparts. Especially interesting is the fact that the largest QTL for three morphological traits critical to pollination success are all within a few centimorgans of each other and have matching divergence directions: spur length is at 24.6cM, pistil length at 32.6cM, and spur curvature at 36cM on chr7.

Nearly all of the morphological traits mapped in this study are controlled by multiple loci of small- and medium-effect, in keeping with the pattern established by previous work in other floral systems in which complex developmental traits have equally complex genetic architectures (Goodwillie et al. 2006; Nakazato et al. 2013; Wessinger et al. 2014). One trait, spur curvature, broke from this norm, with a single locus of major effect on chr7 (as well as two very minor ones on chr 2 & 4). This supports earlier work by Prazmo (1965), which showed that F2 hybrids of *A. canadensis* and *A. flabellata* exhibit a 3:1 ratio of curved:straight spurs, indicating the presence of just one causative locus. Spur curvature is not the only example of a single locus of major effect underlying a morphological trait in *Aquilegia:* the presence of the petal nectar spur itself maps to a single locus *POPOVICH*, which encodes a C2H2-zinc finger transcription factor (Ballerini et al. 2020). Single large loci controlling floral morphological variation remain rare in the QTL mapping literature, such as *L02*, an HLH transcription factor responsible for style length in *Solanum* (Chen et al. 2007), as well as a stigma exertion locus in *Oryza* (Miyata et al. 2007). However, examples of large-effect loci acting in concert with several smaller loci to shape floral organs are quite common. This pattern is found in the QTL maps of sepal length in *Iris* (Bouck et al. 2007); sepal length, petal width, and pistil length in *Arabidopsis* (Juenger et al. 2005); and multiple floral organ size traits in *Petunia* (Hermann et al. 2015).

In this study we characterized the genetic architecture of 17 floral traits encompassing color, nectar composition, and organ morphology that shifted during a bee-to-hummingbird pollinator transition in sister species of *Aquilegia*. This work represents the first comprehensive QTL analysis of *Aquilegia* pollination syndromes using genomic data, and will lead to exciting future work such as validating the ABP model, exploring the numerous CBP candidate genes, and characterizing the developmental basis of morphological traits in an effort to clarify potential causative loci. Given that there is a second independent transition to hummingbird pollination in the genus, and the multiple independent gains and losses of carotenoids, opportunities for comparative studies abound. When we synthesize our findings in *Aquilegia* with previous QTL studies in other systems, we see that every kind of genetic architecture is possible during rapid radiations. This study alone shows that color can be influenced by many loci of small- and moderate-effect, while nectar spur shape can be sculpted by a single locus of large-effect: perhaps it is time to retire the notion of a dichotomy between genetic architectures of biochemical traits and morphological traits.

## Supporting information

Supplemental figures, Table 1, bibliography

## Author contributions

ESB conceived of the study, and designed it with MBE. ESB grew, crossed, and sequenced the parental species, and grew and pollinated the F1 individuals. MBE and YM grew the F2s. GPTC and LM developed the spur curvature phenotyping method, which also yielded spur length phenotypes. AD phenotyped nectary area. All other phenotyping was conducted by MBE. ESB, MBE, and YM prepared the F2 sequencing libraries and constructed the genetic map. MBE and ESB performed QTL mapping analyses, with assistance from AD for nectary area. ND identified anthocyanin and carotenoid loci, and developed the anthocyanin model with ESB and MBE. EMK and SAH provided oversight of the study. MBE wrote the manuscript with input from the co-authors.

## Acknowledgements

We would like to thank the Arnold Arboretum of Harvard University for their generous accommodation of our plants in their Weld Hill growth facilities and use of lab space while phenotyping; Kea Woodruff and Laura Craig-Comin for their plant pest management expertise; Olivia Meyerson and Michael Shahandeh for their discussions of QTL analysis; Joanna Ladopoulu for her assistance with phenotyping; Natasha Parikh for her guidance with Matlab; Elizabeth McCarthy for her advice about floral color; Karl Broman for answering questions about the R/qtl package and maintaining his Google group on the subject; and the WAWG for their thoughtful manuscript suggestions. This material is based upon work supported by the National Science Foundation Graduate Research Fellowship under grant no. DGE1745303 (to MBE). We acknowledge the Harvard Quantitative Biology Initiative and the NSF-Simons Center for Mathematical and Statistical Analysis of Biology at Harvard, award no. 1764269 (to GPTC and LM). ESB was supported by the NIH under the Ruth L. Kirschstein National Research Service Award (F32GM103154). Research was funded by the UC Santa Barbara Harvey Karp Discovery award to ESB. The sequencing was carried out by the DNA Technologies and Expression Analysis Cores at the UC Davis Genome Center, supported by NIH Shared Instrumentation Grant 1S10OD010786-01 and the Biological Nanostructures Lab at UC Santa Barbara. We thank Jennifer Smith and the use of the research facilities within the California NanoSystems Institute, supported by the University of California, Santa Barbara and the University of California, Office of the President.

## Data Accessibility Statement

All sequence data are deposited in the Sequence Read Archive under BioProject ID PRJNA720109.

