## Supplemental figures, Table 1, bibliography for "Genetic architecture of floral traits in bee- and hummingbird-pollinated sister species of *Aquilegia* (columbine)"

This file includes:

Figs. S1-S8
Table S1 (Tables S2-S4 are in separate .xlsx files)
Supplementary bibliography

Figure S1A

PAL

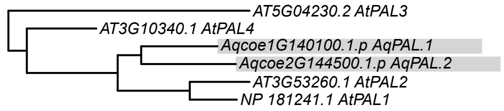


C4H

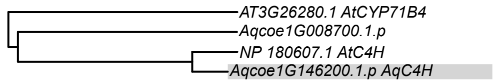


4CL

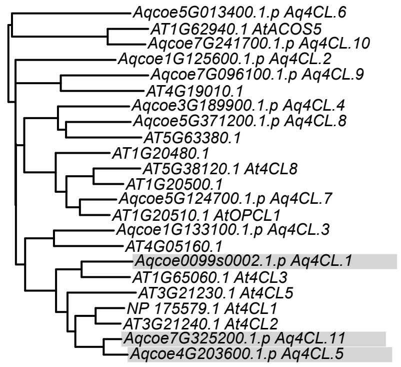

### Figure S1B

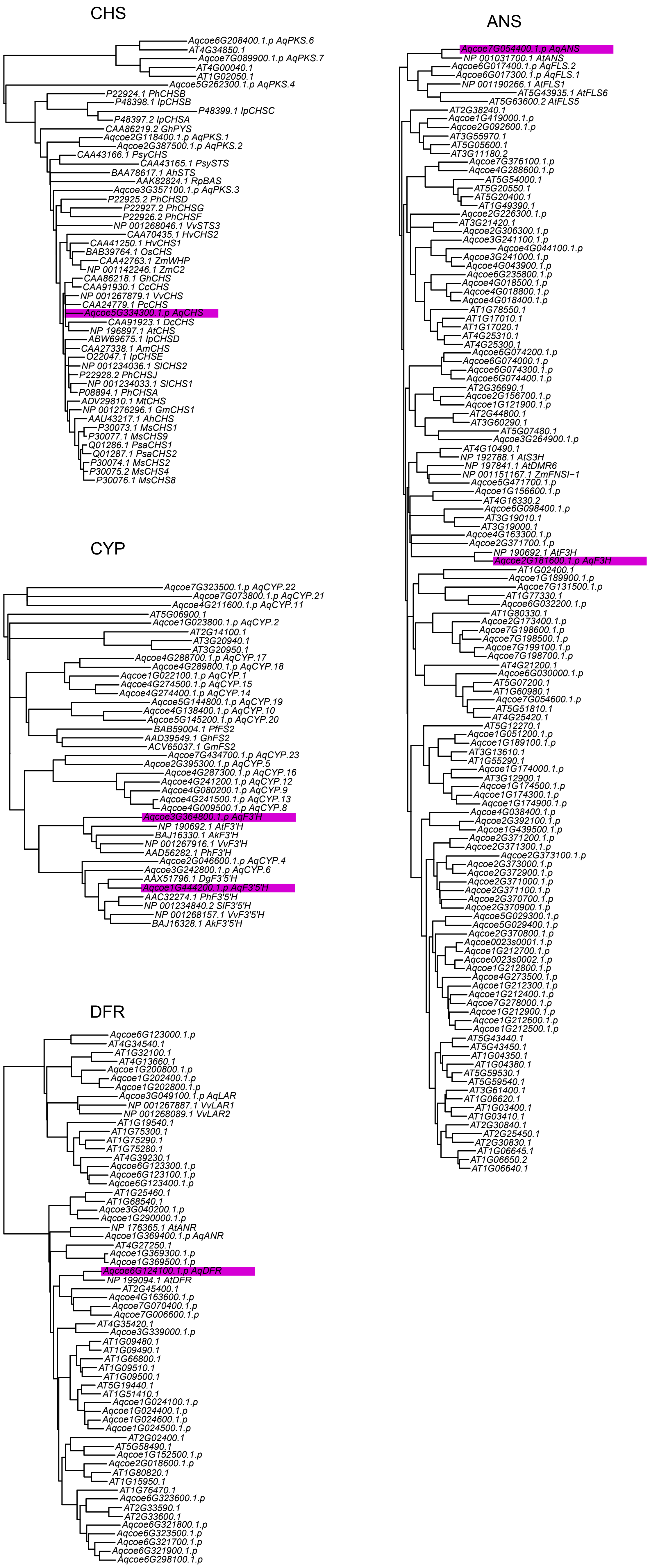

CHS
Aqcoe6G208400.1.p AqPKS.6
AT4G34850.1
Aqcoe7G089900.1.p AqPKS.7
AT4G00040.1
AT1G02050.1
Aqcoe5G262300.1.p AqPKS.4
P22924.1 PhCHSB
P48398.1 IpCHSB
P48399.1 IpCHSC
P48397.2 IpCHSA
CAA86219.2 GhPYS
Aqcoe2G118400.1.p AqPKS.1
Aqcoe2G387500.1.p AqPKS.2
CAA43166.1 PsyCHS
CAA43165.1 PsySTS
BAA78617.1 AhSTS
AAK82824.1 RpBAS
Aqcoe3G357100.1.p AqPKS.3
P22925.2 PhCHSD
P22927.2 PhCHSG
P22926.2 PhCHSF
NP 001268046.1 VvSTS3
CAA70435.1 HvCHS2
CAA41250.1 HvCHS1
BAB39764.1 OsCHS
CAA42763.1 ZmWHP
NP 001142246.1 ZmC2
CAA86218.1 GhCHS
CAA91930.1 CcCHS
NP 001267879.1 VvCHS
CAA24779.1 PcCHS
Aqcoe5G334300.1.p AqCHS
CAA91923.1 DcCHS
NP 196897.1 AtCHS
ABW69675.1 IpCHSD
CAA27338.1 AmCHS
O22047.1 IpCHSE
NP 001234036.1 SlCHS2
P22928.2 PhCHSJ
NP 001234033.1 SlCHS1
P08894.1 PhCHSA
ADV29810.1 MtCHS
NP 001276296.1 GmCHS1
AAU43217.1 AhCHS
P30073.1 MsCHS1
P30077.1 MsCHS9
Q01286.1 PsaCHS1
Q01287.1 PsaCHS2
P30074.1 MsCHS2
P30075.2 MsCHS4
P30076.1 MsCHS8
CYP
Aqcoe7G323500.1.p AqCYP.22
Aqcoe7G073800.1.p AqCYP.21
Aqcoe4G211600.1.p AqCYP.11
AT5G06900.1
Aqcoe1G023800.1.p AqCYP.2
AT2G14100.1
AT3G20940.1
AT3G20950.1
Aqcoe4G288700.1.p AqCYP.17
Aqcoe4G289800.1.p AqCYP.18
Aqcoe1G022100.1.p AqCYP.1
Aqcoe4G274500.1.p AqCYP.15
Aqcoe4G274400.1.p AqCYP.14
Aqcoe5G144800.1.p AqCYP.19
Aqcoe4G138400.1.p AqCYP.10
Aqcoe5G145200.1.p AqCYP.20
BAB59004.1 PfFS2
AAD39549.1 GhFS2
ACV65037.1 GmFS2
Aqcoe7G434700.1.p AqCYP.23
Aqcoe2G395300.1.p AqCYP.5
Aqcoe4G287300.1.p AqCYP.16
Aqcoe4G241200.1.p AqCYP.12
Aqcoe4G080200.1.p AqCYP.9
Aqcoe4G241500.1.p AqCYP.13
Aqcoe4G009500.1.p AqCYP.8
Aqcoe3G364800.1.p AqF3'H
NP 190692.1 AtF3'H
BAJ16330.1 AkF3'H
NP 001267916.1 VvF3'H
AAD56282.1 PhF3'H
Aqcoe2G046600.1.p AqCYP.4
Aqcoe3G242800.1.p AqCYP.6
AAX51796.1 DgF3'5'H
Aqcoe1G444200.1.p AqF3'5'H
AAC32274.1 PhF3'5'H
NP 001234840.2 SlF3'5'H
NP 001268157.1 VvF3'5'H
BAJ16328.1 AkF3'5'H
DFR
Aqcoe6G123000.1.p
AT4G34540.1
AT1G32100.1
AT4G13660.1
Aqcoe1G200800.1.p
Aqcoe1G202400.1.p
Aqcoe1G202800.1.p
Aqcoe3G049100.1.p AqLAR
NP 001267887.1 VvLAR1
NP 001268089.1 VvLAR2
AT1G19540.1
AT1G75300.1
AT1G75290.1
AT1G75280.1
AT4G39230.1
Aqcoe6G123300.1.p
Aqcoe6G123100.1.p
Aqcoe6G123400.1.p
AT1G25460.1
AT1G68540.1
Aqcoe3G040200.1.p
Aqcoe1G290000.1.p
NP 176365.1 AtANR
Aqcoe1G369400.1.p AqANR
AT4G27250.1
Aqcoe1G369300.1.p
Aqcoe1G369500.1.p
Aqcoe6G124100.1.p AqDFR
NP 199094.1 AtDFR
AT2G45400.1
Aqcoe4G163600.1.p
Aqcoe7G070400.1.p
Aqcoe7G006600.1.p
AT4G35420.1
Aqcoe3G339000.1.p
AT1G09480.1
AT1G09490.1
AT1G66800.1
AT1G09510.1
AT1G09500.1
AT5G19440.1
AT1G51410.1
Aqcoe1G024100.1.p
Aqcoe1G024400.1.p
Aqcoe1G024600.1.p
Aqcoe1G024500.1.p
AT2G02400.1
AT5G58490.1
Aqcoe1G152500.1.p
Aqcoe2G018600.1.p
AT1G80820.1
AT1G15950.1
AT1G76470.1
Aqcoe6G323600.1.p
AT2G33590.1
AT2G33600.1
Aqcoe6G321800.1.p
Aqcoe6G323500.1.p
Aqcoe6G321700.1.p
Aqcoe6G321900.1.p
Aqcoe6G298100.1.p
ANS
Aqcoe7G054400.1.p AqANS
NP 001031700.1 AtANS
Aqcoe6G017400.1.p AqFLS.2
Aqcoe6G017300.1.p AqFLS.1
NP 001190266.1 AtFLS1
AT5G43935.1 AtFLS6
AT5G63600.2 AtFLS5
AT2G38240.1
Aqcoe1G419000.1.p
Aqcoe2G092600.1.p
AT3G55970.1
AT5G05600.1
AT3G11180.2
Aqcoe7G376100.1.p
Aqcoe4G288600.1.p
AT5G54000.1
AT5G20550.1
AT5G20400.1
AT1G49390.1
Aqcoe2G226300.1.p
AT3G21420.1
Aqcoe2G306300.1.p
Aqcoe3G241100.1.p
Aqcoe4G044100.1.p
Aqcoe3G241000.1.p
Aqcoe4G043900.1.p
Aqcoe6G235800.1.p
Aqcoe4G018500.1.p
Aqcoe4G018800.1.p
Aqcoe4G018400.1.p
AT1G78550.1
AT1G17010.1
AT1G17020.1
AT4G25310.1
AT4G25300.1
Aqcoe6G074200.1.p
Aqcoe6G074000.1.p
Aqcoe6G074300.1.p
Aqcoe6G074400.1.p
AT2G36690.1
Aqcoe2G156700.1.p
Aqcoe1G121900.1.p
AT2G44800.1
AT3G60290.1
AT5G07480.1
Aqcoe3G264900.1.p
AT4G10490.1
NP 192788.1 AtS3H
NP 197841.1 AtDMR6
NP 001151167.1 ZmFNSI−1
Aqcoe5G471700.1.p
Aqcoe1G156600.1.p
AT4G16330.2
Aqcoe6G098400.1.p
AT3G19010.1
AT3G19000.1
Aqcoe4G163300.1.p
Aqcoe2G371700.1.p
NP 190692.1 AtF3H
Aqcoe2G181600.1.p AqF3H
AT1G02400.1
Aqcoe1G189900.1.p
Aqcoe7G131500.1.p
AT1G77330.1
Aqcoe6G032200.1.p
AT1G80330.1
Aqcoe2G173400.1.p
Aqcoe7G198600.1.p
Aqcoe7G198500.1.p
Aqcoe7G199100.1.p
Aqcoe7G198700.1.p
AT4G21200.1
Aqcoe6G030000.1.p
AT5G07200.1
AT1G60980.1
Aqcoe7G054600.1.p
AT5G51810.1
AT4G25420.1
AT5G12270.1
Aqcoe1G051200.1.p
Aqcoe1G189100.1.p
AT3G13610.1
AT1G55290.1
Aqcoe1G174000.1.p
AT3G12900.1
Aqcoe1G174500.1.p
Aqcoe1G174300.1.p
Aqcoe1G174900.1.p
Aqcoe4G038400.1.p
Aqcoe2G392100.1.p
Aqcoe1G439500.1.p
Aqcoe2G371200.1.p
Aqcoe2G371300.1.p
Aqcoe2G373100.1.p
Aqcoe2G373000.1.p
Aqcoe2G372900.1.p
Aqcoe2G371000.1.p
Aqcoe2G371100.1.p
Aqcoe2G370700.1.p
Aqcoe2G370900.1.p
Aqcoe5G029300.1.p
Aqcoe5G029400.1.p
Aqcoe2G370800.1.p
Aqcoe0023s0001.1.p
Aqcoe1G212700.1.p
Aqcoe0023s0002.1.p
Aqcoe1G212800.1.p
Aqcoe4G273500.1.p
Aqcoe1G212300.1.p
Aqcoe1G212400.1.p
Aqcoe7G278000.1.p
Aqcoe1G212900.1.p
Aqcoe1G212600.1.p
Aqcoe1G212500.1.p
AT5G43440.1
AT5G43450.1
AT1G04350.1
AT1G04380.1
AT5G59530.1
AT5G59540.1
AT3G61400.1
AT1G06620.1
AT1G03400.1
AT1G03410.1
AT2G30840.1
AT2G25450.1
AT2G30830.1
AT1G06645.1
AT1G06650.2
AT1G06640.1

### Figure S1C

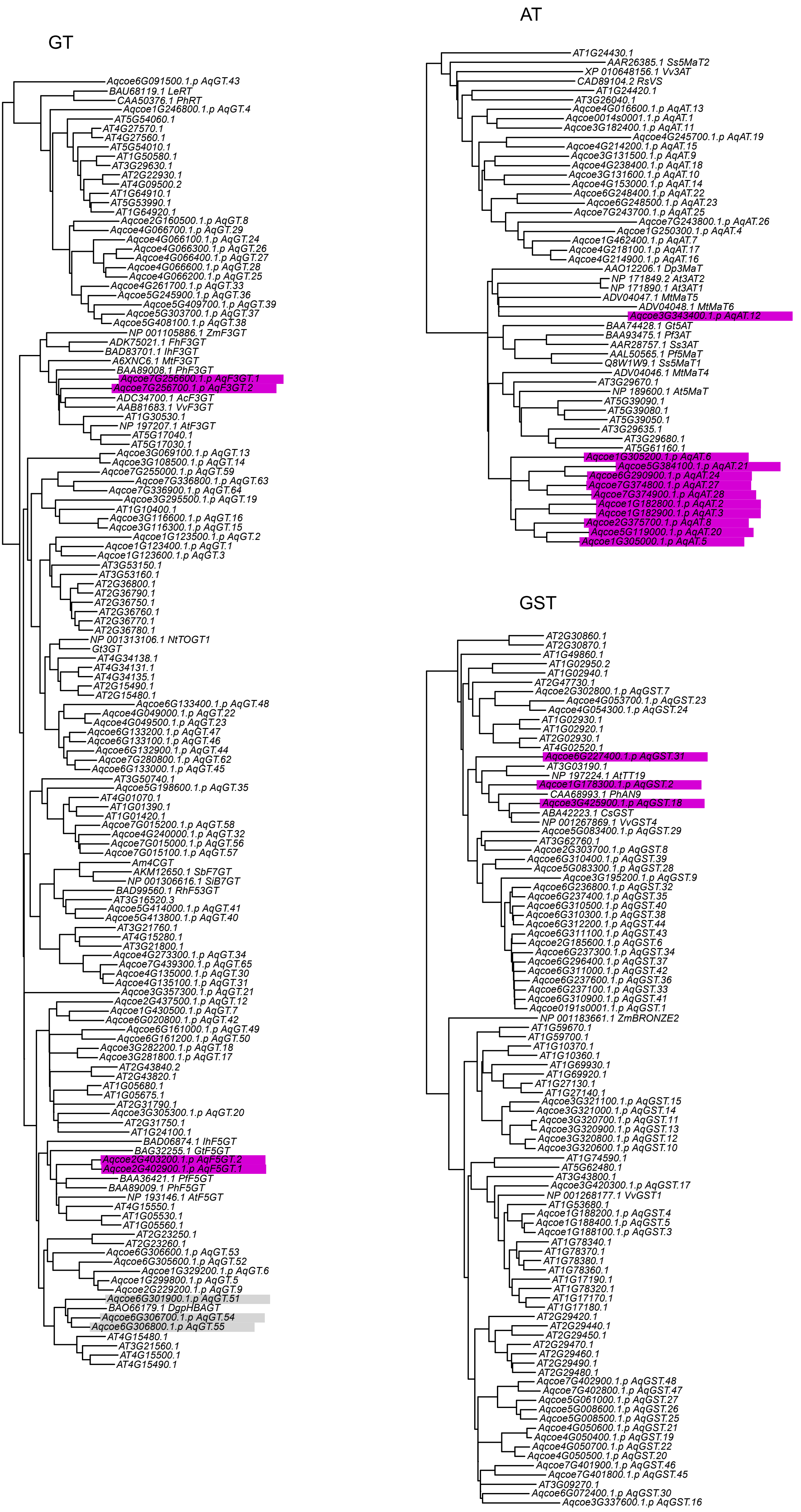
GT
Aqcoe6G091500.1.p AqGT.43
BAU68119.1 LeRT
CAA50376.1 PhRT
Aqcoe1G246800.1.p AqGT.4
AT5G54060.1
AT4G27570.1
AT4G27560.1
AT5G54010.1
AT1G50580.1
AT3G29630.1
AT2G22930.1
AT4G09500.2
AT1G64910.1
AT5G53990.1
AT1G64920.1
Aqcoe2G160500.1.p AqGT.8
Aqcoe4G066700.1.p AqGT.29
Aqcoe4G066100.1.p AqGT.24
Aqcoe4G066300.1.p AqGT.26
Aqcoe4G066400.1.p AqGT.27
Aqcoe4G066600.1.p AqGT.28
Aqcoe4G066200.1.p AqGT.25
Aqcoe4G261700.1.p AqGT.33
Aqcoe5G245900.1.p AqGT.36
Aqcoe5G409700.1.p AqGT.39
Aqcoe5G303700.1.p AqGT.37
Aqcoe5G408100.1.p AqGT.38
NP 001105886.1 ZmF3GT
ADK75021.1 FhF3GT
BAD83701.1 IhF3GT
A6XNC6.1 MtF3GT
BAA89008.1 PhF3GT
Aqcoe7G256600.1.p AqF3GT.1
Aqcoe7G256700.1.p AqF3GT.2
ADC34700.1 AcF3GT
AAB81683.1 VvF3GT
AT1G30530.1
NP 197207.1 AtF3GT
AT5G17040.1
AT5G17030.1
Aqcoe3G069100.1.p AqGT.13
Aqcoe3G108500.1.p AqGT.14
Aqcoe7G255000.1.p AqGT.59
Aqcoe7G336800.1.p AqGT.63
Aqcoe7G336900.1.p AqGT.64
Aqcoe3G295500.1.p AqGT.19
AT1G10400.1
Aqcoe3G116600.1.p AqGT.16
Aqcoe3G116300.1.p AqGT.15
Aqcoe1G123500.1.p AqGT.2
Aqcoe1G123400.1.p AqGT.1
Aqcoe1G123600.1.p AqGT.3
AT3G53150.1
AT3G53160.1
AT2G36800.1
AT2G36790.1
AT2G36750.1
AT2G36760.1
AT2G36770.1
AT2G36780.1
NP 001313106.1 NtTOGT1
Gt3GT
AT4G34138.1
AT4G34131.1
AT4G34135.1
AT2G15490.1
AT2G15480.1
Aqcoe6G133400.1.p AqGT.48
Aqcoe4G049000.1.p AqGT.22
Aqcoe4G049500.1.p AqGT.23
Aqcoe6G133200.1.p AqGT.47
Aqcoe6G133100.1.p AqGT.46
Aqcoe6G132900.1.p AqGT.44
Aqcoe7G280800.1.p AqGT.62
Aqcoe6G133000.1.p AqGT.45
AT3G50740.1
Aqcoe5G198600.1.p AqGT.35
AT4G01070.1
AT1G01390.1
AT1G01420.1
Aqcoe7G015200.1.p AqGT.58
Aqcoe4G240000.1.p AqGT.32
Aqcoe7G015000.1.p AqGT.56
Aqcoe7G015100.1.p AqGT.57
Am4CGT
AKM12650.1 SbF7GT
NP 001306616.1 SiB7GT
BAD99560.1 RhF53GT
AT3G16520.3
Aqcoe5G414000.1.p AqGT.41
Aqcoe5G413800.1.p AqGT.40
AT3G21760.1
AT4G15280.1
AT3G21800.1
Aqcoe4G273300.1.p AqGT.34
Aqcoe7G439300.1.p AqGT.65
Aqcoe4G135000.1.p AqGT.30
Aqcoe4G135100.1.p AqGT.31
Aqcoe3G357300.1.p AqGT.21
Aqcoe2G437500.1.p AqGT.12
Aqcoe1G430500.1.p AqGT.7
Aqcoe6G020800.1.p AqGT.42
Aqcoe6G161000.1.p AqGT.49
Aqcoe6G161200.1.p AqGT.50
Aqcoe3G282200.1.p AqGT.18
Aqcoe3G281800.1.p AqGT.17
AT2G43840.2
AT2G43820.1
AT1G05680.1
AT1G05675.1
AT2G31790.1
Aqcoe3G305300.1.p AqGT.20
AT2G31750.1
AT1G24100.1
BAD06874.1 IhF5GT
BAG32255.1 GtF5GT
Aqcoe2G403200.1.p AqF5GT.2
Aqcoe2G402900.1.p AqF5GT.1
BAA36421.1 PfF5GT
BAA89009.1 PhF5GT
NP 193146.1 AtF5GT
AT4G15550.1
AT1G05530.1
AT1G05560.1
AT2G23250.1
AT2G23260.1
Aqcoe6G306600.1.p AqGT.53
Aqcoe6G305600.1.p AqGT.52
Aqcoe1G329200.1.p AqGT.6
Aqcoe1G299800.1.p AqGT.5
Aqcoe2G229200.1.p AqGT.9
Aqcoe6G301900.1.p AqGT.51
BAO66179.1 DgpHBAGT
Aqcoe6G306700.1.p AqGT.54
Aqcoe6G306800.1.p AqGT.55
AT4G15480.1
AT3G21560.1
AT4G15500.1
AT4G15490.1
AT
AT1G24430.1
AAR26385.1 Ss5MaT2
XP 010648156.1 Vv3AT
CAD89104.2 RsVS
AT1G24420.1
AT3G26040.1
Aqcoe4G016600.1.p AqAT.13
Aqcoe0014s0001.1.p AqAT.1
Aqcoe3G182400.1.p AqAT.11
Aqcoe4G245700.1.p AqAT.19
Aqcoe4G214200.1.p AqAT.15
Aqcoe3G131500.1.p AqAT.9
Aqcoe4G238400.1.p AqAT.18
Aqcoe3G131600.1.p AqAT.10
Aqcoe4G153000.1.p AqAT.14
Aqcoe6G248400.1.p AqAT.22
Aqcoe6G248500.1.p AqAT.23
Aqcoe7G243700.1.p AqAT.25
Aqcoe7G243800.1.p AqAT.26
Aqcoe1G250300.1.p AqAT.4
Aqcoe1G462400.1.p AqAT.7
Aqcoe4G218100.1.p AqAT.17
Aqcoe4G214900.1.p AqAT.16
AAO12206.1 Dp3MaT
NP 171849.2 At3AT2
NP 171890.1 At3AT1
ADV04047.1 MtMaT5
ADV04048.1 MtMaT6
Aqcoe3G343400.1.p AqAT.12
BAA74428.1 Gt5AT
BAA93475.1 Pf3AT
AAR28757.1 Ss3AT
AAL50565.1 Pf5MaT
Q8W1W9.1 Ss5MaT1
ADV04046.1 MtMaT4
AT3G29670.1
NP 189600.1 At5MaT
AT5G39090.1
AT5G39080.1
AT5G39050.1
AT3G29635.1
AT3G29680.1
AT5G61160.1
Aqcoe1G305200.1.p AqAT.6
Aqcoe5G384100.1.p AqAT.21
Aqcoe6G290900.1.p AqAT.24
Aqcoe7G374800.1.p AqAT.27
Aqcoe7G374900.1.p AqAT.28
Aqcoe1G182800.1.p AqAT.2
Aqcoe1G182900.1.p AqAT.3
Aqcoe2G375700.1.p AqAT.8
Aqcoe5G119000.1.p AqAT.20
Aqcoe1G305000.1.p AqAT.5
GST
AT2G30860.1
AT2G30870.1
AT1G49860.1
AT1G02950.2
AT1G02940.1
AT2G47730.1
Aqcoe2G302800.1.p AqGST.7
Aqcoe4G053700.1.p AqGST.23
Aqcoe4G054300.1.p AqGST.24
AT1G02930.1
AT1G02920.1
AT2G02930.1
AT4G02520.1
Aqcoe6G227400.1.p AqGST.31
AT3G03190.1
NP 197224.1 AtTT19
Aqcoe1G178300.1.p AqGST.2
CAA68993.1 PhAN9
Aqcoe3G425900.1.p AqGST.18
ABA42223.1 CsGST
NP 001267869.1 VvGST4
Aqcoe5G083400.1.p AqGST.29
AT3G62760.1
Aqcoe2G303700.1.p AqGST.8
Aqcoe6G310400.1.p AqGST.39
Aqcoe5G083300.1.p AqGST.28
Aqcoe3G195200.1.p AqGST.9
Aqcoe6G236800.1.p AqGST.32
Aqcoe6G237400.1.p AqGST.35
Aqcoe6G310500.1.p AqGST.40
Aqcoe6G310300.1.p AqGST.38
Aqcoe6G312200.1.p AqGST.44
Aqcoe6G311100.1.p AqGST.43
Aqcoe2G185600.1.p AqGST.6
Aqcoe6G237300.1.p AqGST.34
Aqcoe6G296400.1.p AqGST.37
Aqcoe6G311000.1.p AqGST.42
Aqcoe6G237600.1.p AqGST.36
Aqcoe6G237100.1.p AqGST.33
Aqcoe6G310900.1.p AqGST.41
Aqcoe0191s0001.1.p AqGST.1
NP 001183661.1 ZmBRONZE2
AT1G59670.1
AT1G59700.1
AT1G10370.1
AT1G10360.1
AT1G69930.1
AT1G69920.1
AT1G27130.1
AT1G27140.1
Aqcoe3G321100.1.p AqGST.15
Aqcoe3G321000.1.p AqGST.14
Aqcoe3G320700.1.p AqGST.11
Aqcoe3G320900.1.p AqGST.13
Aqcoe3G320800.1.p AqGST.12
Aqcoe3G320600.1.p AqGST.10
AT1G74590.1
AT5G62480.1
AT3G43800.1
Aqcoe3G420300.1.p AqGST.17
NP 001268177.1 VvGST1
AT1G53680.1
Aqcoe1G188200.1.p AqGST.4
Aqcoe1G188400.1.p AqGST.5
Aqcoe1G188100.1.p AqGST.3
AT1G78340.1
AT1G78370.1
AT1G78380.1
AT1G78360.1
AT1G17190.1
AT1G78320.1
AT1G17170.1
AT1G17180.1
AT2G29420.1
AT2G29440.1
AT2G29450.1
AT2G29470.1
AT2G29460.1
AT2G29490.1
AT2G29480.1
Aqcoe7G402900.1.p AqGST.48
Aqcoe7G402800.1.p AqGST.47
Aqcoe5G061000.1.p AqGST.27
Aqcoe5G008600.1.p AqGST.26
Aqcoe5G008500.1.p AqGST.25
Aqcoe4G050600.1.p AqGST.21
Aqcoe4G050400.1.p AqGST.19
Aqcoe4G050700.1.p AqGST.22
Aqcoe4G050500.1.p AqGST.20
Aqcoe7G401900.1.p AqGST.46
Aqcoe7G401800.1.p AqGST.45
AT3G09270.1
Aqcoe6G072400.1.p AqGST.30
Aqcoe3G337600.1.p AqGST.16

Figure S1D

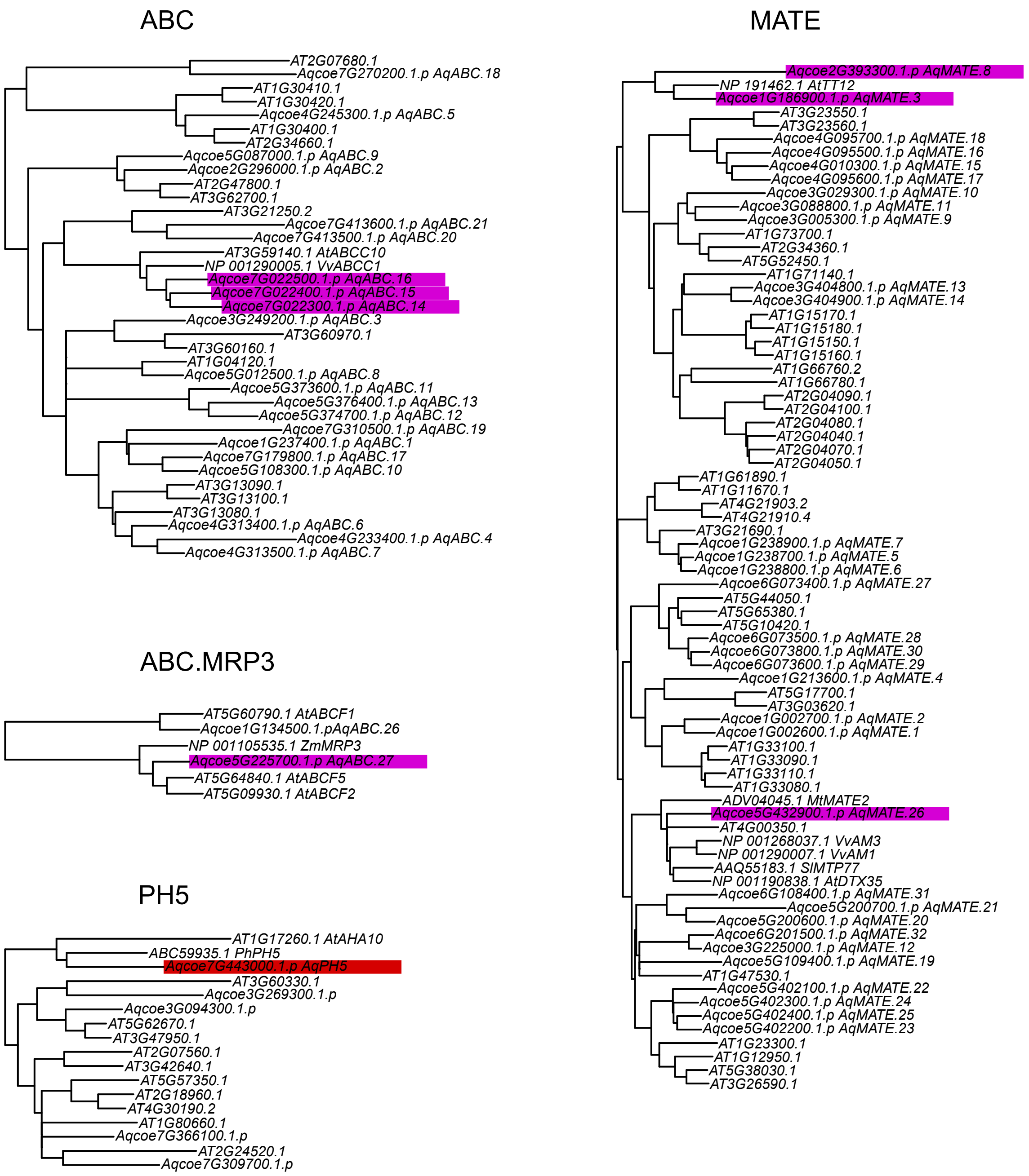
ABC
AT2G07680.1
Aqcoe7G270200.1.p AqABC.18
AT1G30410.1
AT1G30420.1
Aqcoe4G245300.1.p AqABC.5
AT1G30400.1
AT2G34660.1
Aqcoe5G087000.1.p AqABC.9
Aqcoe2G296000.1.p AqABC.2
AT2G47800.1
AT3G62700.1
AT3G21250.2
Aqcoe7G413600.1.p AqABC.21
Aqcoe7G413500.1.p AqABC.20
AT3G59140.1 AtABCC10
NP 001290005.1 VvABCC1
Aqcoe7G022500.1.p AqABC.16
Aqcoe7G022400.1.p AqABC.15
Aqcoe7G022300.1.p AqABC.14
Aqcoe3G249200.1.p AqABC.3
AT3G60970.1
AT3G60160.1
AT1G04120.1
Aqcoe5G012500.1.p AqABC.8
Aqcoe5G373600.1.p AqABC.11
Aqcoe5G376400.1.p AqABC.13
Aqcoe5G374700.1.p AqABC.12
Aqcoe7G310500.1.p AqABC.19
Aqcoe1G237400.1.p AqABC.1
Aqcoe7G179800.1.p AqABC.17
Aqcoe5G108300.1.p AqABC.10
AT3G13090.1
AT3G13100.1
AT3G13080.1
Aqcoe4G313400.1.p AqABC.6
Aqcoe4G233400.1.p AqABC.4
Aqcoe4G313500.1.p AqABC.7
ABC.MRP3
AT5G60790.1 AtABCF1
Aqcoe1G134500.1.pAqABC.26
NP 001105535.1 ZmMRP3
Aqcoe5G225700.1.p AqABC.27
AT5G64840.1 AtABCF5
AT5G09930.1 AtABCF2
PH5
AT1G17260.1 AtAHA10
ABC59935.1 PhPH5
Aqcoe7G443000.1.p AqPH5
AT3G60330.1
Aqcoe3G269300.1.p
Aqcoe3G094300.1.p
AT5G62670.1
AT3G47950.1
AT2G07560.1
AT3G42640.1
AT5G57350.1
AT2G18960.1
AT4G30190.2
AT1G80660.1
Aqcoe7G366100.1.p
AT2G24520.1
Aqcoe7G309700.1.p
MATE
Aqcoe2G393300.1.p AqMATE.8
NP 191462.1 AtTT12
Aqcoe1G186900.1.p AqMATE.3
AT3G23550.1
AT3G23560.1
Aqcoe4G095700.1.p AqMATE.18
Aqcoe4G095500.1.p AqMATE.16
Aqcoe4G010300.1.p AqMATE.15
Aqcoe4G095600.1.p AqMATE.17
Aqcoe3G029300.1.p AqMATE.10
Aqcoe3G088800.1.p AqMATE.11
Aqcoe3G005300.1.p AqMATE.9
AT1G73700.1
AT2G34360.1
AT5G52450.1
AT1G71140.1
Aqcoe3G404800.1.p AqMATE.13
Aqcoe3G404900.1.p AqMATE.14
AT1G15170.1
AT1G15180.1
AT1G15150.1
AT1G15160.1
AT1G66760.2
AT1G66780.1
AT2G04090.1
AT2G04100.1
AT2G04080.1
AT2G04040.1
AT2G04070.1
AT2G04050.1
AT1G61890.1
AT1G11670.1
AT4G21903.2
AT4G21910.4
AT3G21690.1
Aqcoe1G238900.1.p AqMATE.7
Aqcoe1G238700.1.p AqMATE.5
Aqcoe1G238800.1.p AqMATE.6
Aqcoe6G073400.1.p AqMATE.27
AT5G44050.1
AT5G65380.1
AT5G10420.1
Aqcoe6G073500.1.p AqMATE.28
Aqcoe6G073800.1.p AqMATE.30
Aqcoe6G073600.1.p AqMATE.29
Aqcoe1G213600.1.p AqMATE.4
AT5G17700.1
AT3G03620.1
Aqcoe1G002700.1.p AqMATE.2
Aqcoe1G002600.1.p AqMATE.1
AT1G33100.1
AT1G33090.1
AT1G33110.1
AT1G33080.1
ADV04045.1 MtMATE2
Aqcoe5G432900.1.p AqMATE.26
AT4G00350.1
NP 001268037.1 VvAM3
NP 001290007.1 VvAM1
AAQ55183.1 SlMTP77
NP 001190838.1 AtDTX35
Aqcoe6G108400.1.p AqMATE.31
Aqcoe5G200700.1.p AqMATE.21
Aqcoe5G200600.1.p AqMATE.20
Aqcoe6G201500.1.p AqMATE.32
Aqcoe3G225000.1.p AqMATE.12
Aqcoe5G109400.1.p AqMATE.19
AT1G47530.1
Aqcoe5G402100.1.p AqMATE.22
Aqcoe5G402300.1.p AqMATE.24
Aqcoe5G402400.1.p AqMATE.25
Aqcoe5G402200.1.p AqMATE.23
AT1G23300.1
AT1G12950.1
AT5G38030.1
AT3G26590.1

### Figure S1E

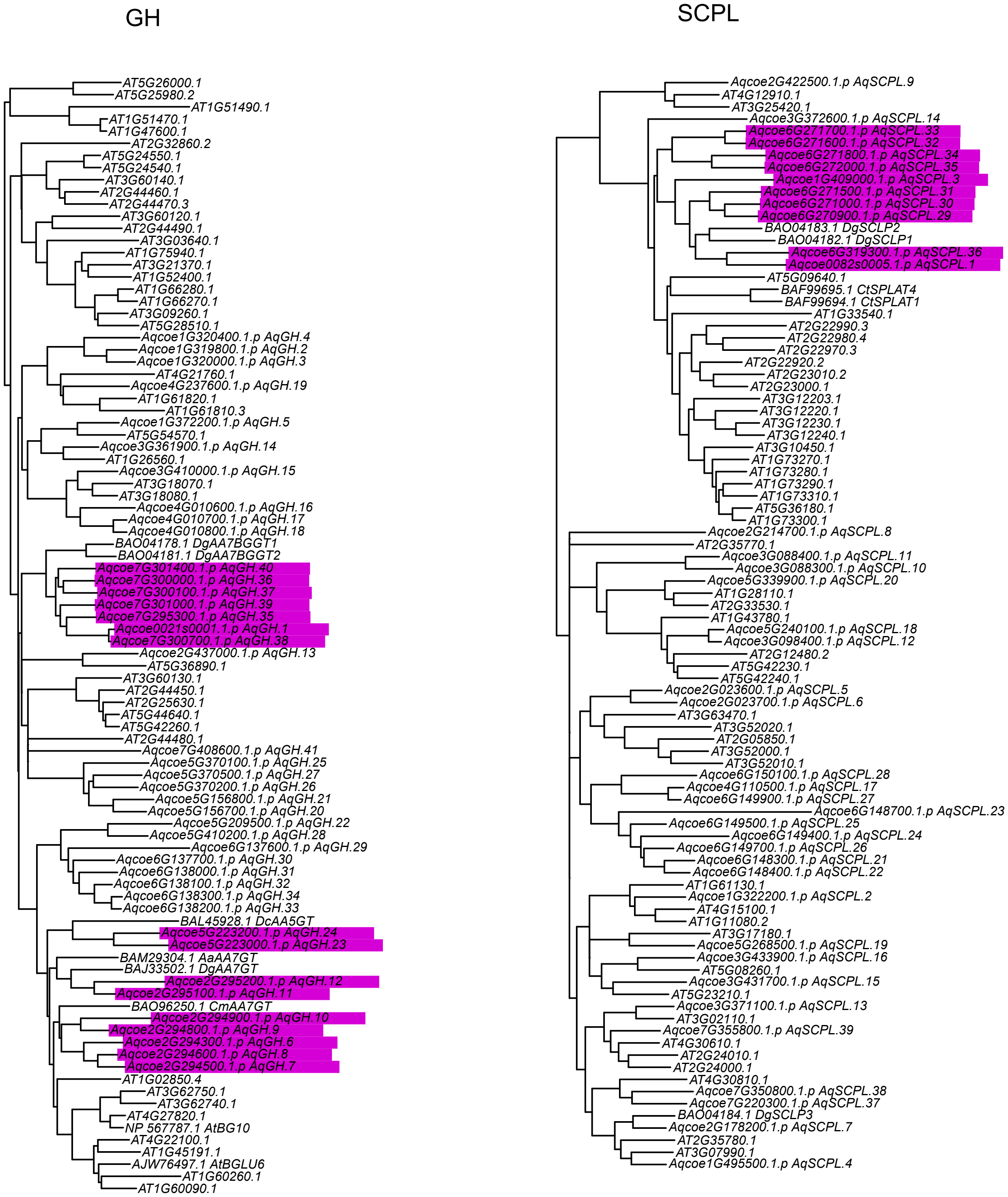

Figure S1F

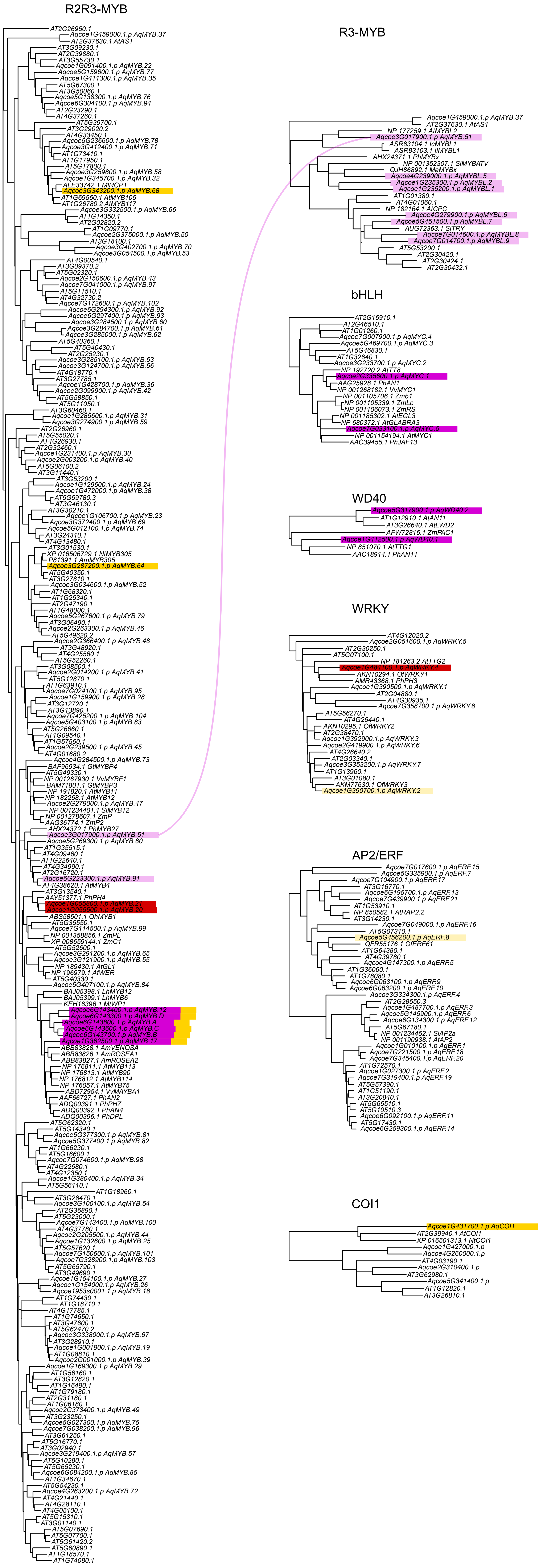

### Figure S1G

DXS

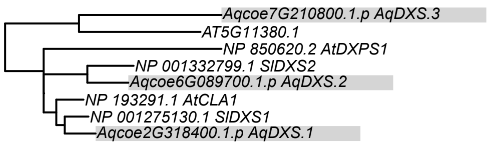


GGPPS

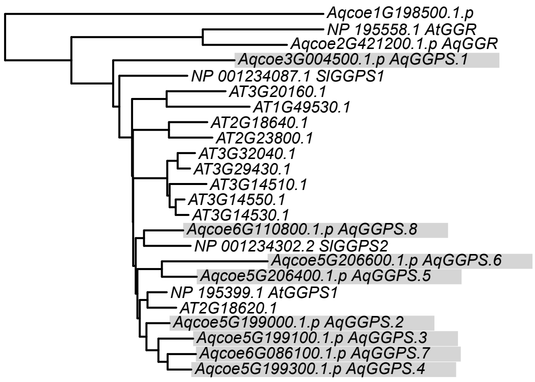

Figure S1H

PSY

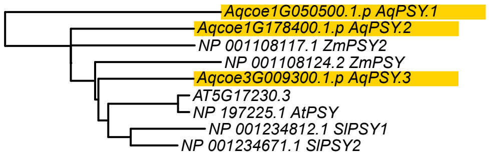


LCYB, LCYE, NSY

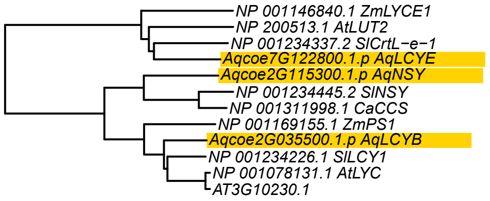


BCH

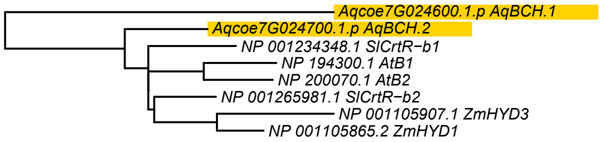


CYP97

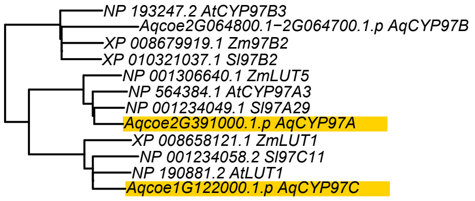


PYP1

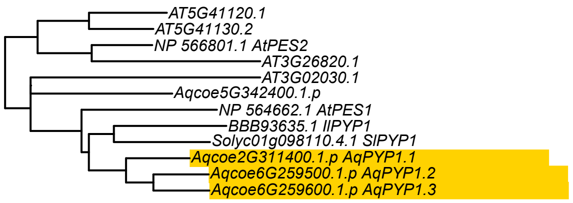

### Figure S1I

CCD/NCED

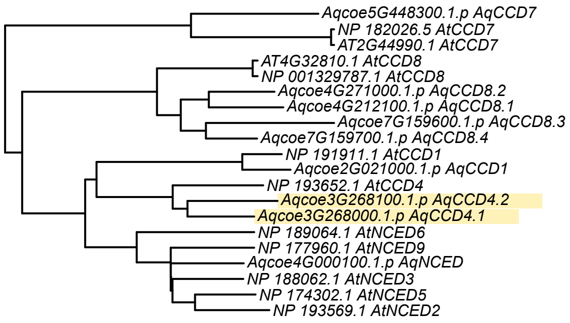


OR

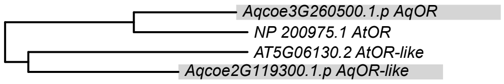


TPRL

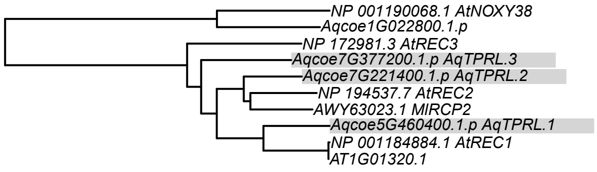

**Figure S1. Neighbor-joining trees of genes involved in anthocyanin and carotenoid pigmentation**

**A.** Neighbor-joining trees of anthocyanin precursor biosynthetic genes. Query group names are abbreviated as: phenylalanine ammonia-lyase (PAL); cinnamate-4-hydroxylase (C4H); and 4-coumaroyl:CoA-ligase (4CL). Likely *Aquilegia* homologs are marked with a gray background.

**B.** Neighbor-joining trees of biosynthetic genes producing the core anthocyanidins pelargonidin, cyanidin, and delphinidin**.** Query group names are abbreviated as: chalcone synthase (CHS); cytochromes P450 (CYP); dihydroflavonol-4-reductase (DFR); and anthocyanidin synthase (ANS). The CYP group included flavonoid 3'-hydroxylases (F3'H) and flavonoid 3',5'-hydroxylases (F3'5'H). The ANS group included ANS and flavanone 3-hydroxylase (F3H). Likely *Aquilegia* homologs are marked with a purple background.

**C.** Neighbor-joining trees of anthocyanidin modification genes**.** Query group names are abbreviated as: glycosyltransferase (GT); acyltransferase (AT); and glutathione S-transferases (GST). GSTs do not appear to modify anthocyanins, but are important for transport into the vacuole (Zhao 2015). The GT group includes anthocyanidin 3-O-glucosyltransferases (F3GT) and anthocyanidin 5-O-glucosyltransferases (F5GT), as well a UDP-glucose dependent p-hydroxybenzoic acid glucosyltransferase (pHBAGT) identified in *Delphinium*. DgpHBAGT plays an indirect role in producing flower color, as its product (pHBG) is a donor molecule for both glycosylation and acylation of anthocyanins in the vacuole (Nishizaki et al 2013). Likely *Aquilegia* homologs of genes directly involved in anthocyanin biosynthesis or transport are marked with a purple background, and homologs of DgpHBAGT are marked with gray.

**D.** Neighbor-joining trees of vacuolar transporter genes. Query group names are abbreviated as: dultidrug resistance-associated protein transporters (ABC, ABC.MRP3); multidrug and toxic compound extrusion transporters (MATE); and H+-APTase (PH5). Likely *Aquilegia* homologs for anthocyanin transporters are marked with a purple background, and the homolog for proton transporter PH5 is marked with red.

**E.** Neighbor-joining trees of anthocyanin modification genes active in the vacuole. Query group names are abbreviated as: glycosyl hydrolase (GH); and serine carboxypeptidase-like proteins (SCPL). Likely *Aquilegia* homologs for anthocyanin transporters are marked with a purple background.

**F.** Neighbor-joining trees of transcriptional regulatory genes control pigment-related genes. Query group names are abbreviated as: myeloblastosis viral oncogene transcription factor (R2R3-MYB, R3-MYB); basic helix-loop-helix transcription factor (bHLH); WD40 repeat containing domain transcription factor (WD40); WRKY transcription factor (WRKY); APETALA2 and ethylene response factor transcription factors (AP2/ERF); and CORONATINE INSENSITIVE1 (COI1). R3-MYBs are found as two separate clades nested within R2R3-MYBs (Gates et al 2018); we pruned the R3-MYB tree to include only those clades plus an outgroup (AtAS1 and Aq.MYB.37). AqMYB5.1 falls within the AtMYBL2 clade, but retains an R2 motif so is classified here as an R2R3-MYB. Transcriptional regulation of floral carotenoid production is poorly characterized. Co-option of an anthocyanin-type R2R3-MYB for control of carotenoid biosynthetic genes has to date only been seen in *Medicago truncatula* (Meng et al 2019), and conserved function of the R2R3-MYB RCP1 from *Erythranthe lewisii* has not yet been demonstrated (Sagawa et al 2016). Similarly, positive regulation of CCD4 by WRKY and ERF transcrption factors is only known from one species, *Osmanthus fragrans* (Han et al 2016, Han et al 2019). Likely *Aquilegia* homologs for transcription factors directly controlling expression anthocyanin biosynthetic genes are marked with a purple background, with light purple indicating negative regulation. *Aquilegia* homologs for transcription factors controlling vacuolar pH are marked with red. Homologs for potential transcriptional regulators of carotenoid biosynthetic genes are marked with yellow, with light yellow indicating positive regulators of CCD4, an enzyme that degrades carotenoids.

**G.** Neighbor-joining trees of carotenoid precursor biosynthetic genes. Query group names are abbreviated as: 1-deoxy-D-xylulose 5-phosphate synthase (DXS); and geranylgeranyl diphosphate synthase (GGPPS). Likely *Aquilegia* homologs are marked with a gray background.

**H.** Neighbor-joining trees of carotenoid biosynthetic and modification genes. Query group names are abbreviated as: phytoene synthase (PSY); lycopene beta-cyclase (LCYB); lycopene epsilon-cyclase (LCYE); neoxanthin synthase (NSY); carotenoid beta-hydroxylase (BCH); and PALE YELLOW PETAL1 (PYP1). Likely *Aquilegia* homologs are marked with a yellow background.

**I.** Neighbor-joining trees of genes indirectly regulating carotenoid biosynthesis and accumulation. Query group names are abbreviated as: carotenoid cleavage dioxygenase (CCD); nine-cis-epoxycarotenoid dioxygenase (NCED); ORANGE protein (OR); and tetratricopeptide repeat-like superfamily protein (TPRL). Likely *Aquilegia* homologs for CCD4 are marked with a pale yellow background, and homolgs for OR and a TPRL from *Erythranthe lewisii* (RCP2) are marked with grey.

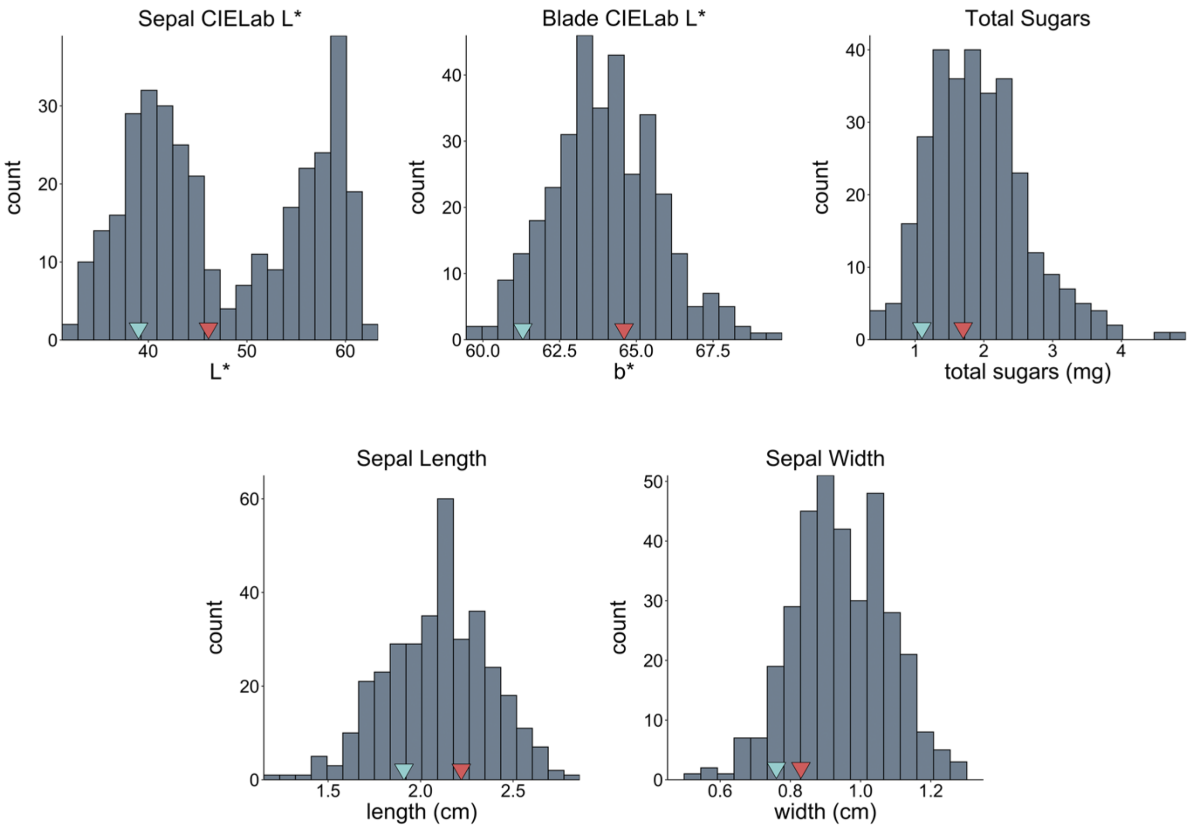


**Figure S2.** F2 population floral trait histograms (continued from Fig. 2). Blue and red arrows mark the phenotypic means of *A. brevistyla* (brev mean) and *A. canadensis* (can mean) plants that were closely related to the parents.

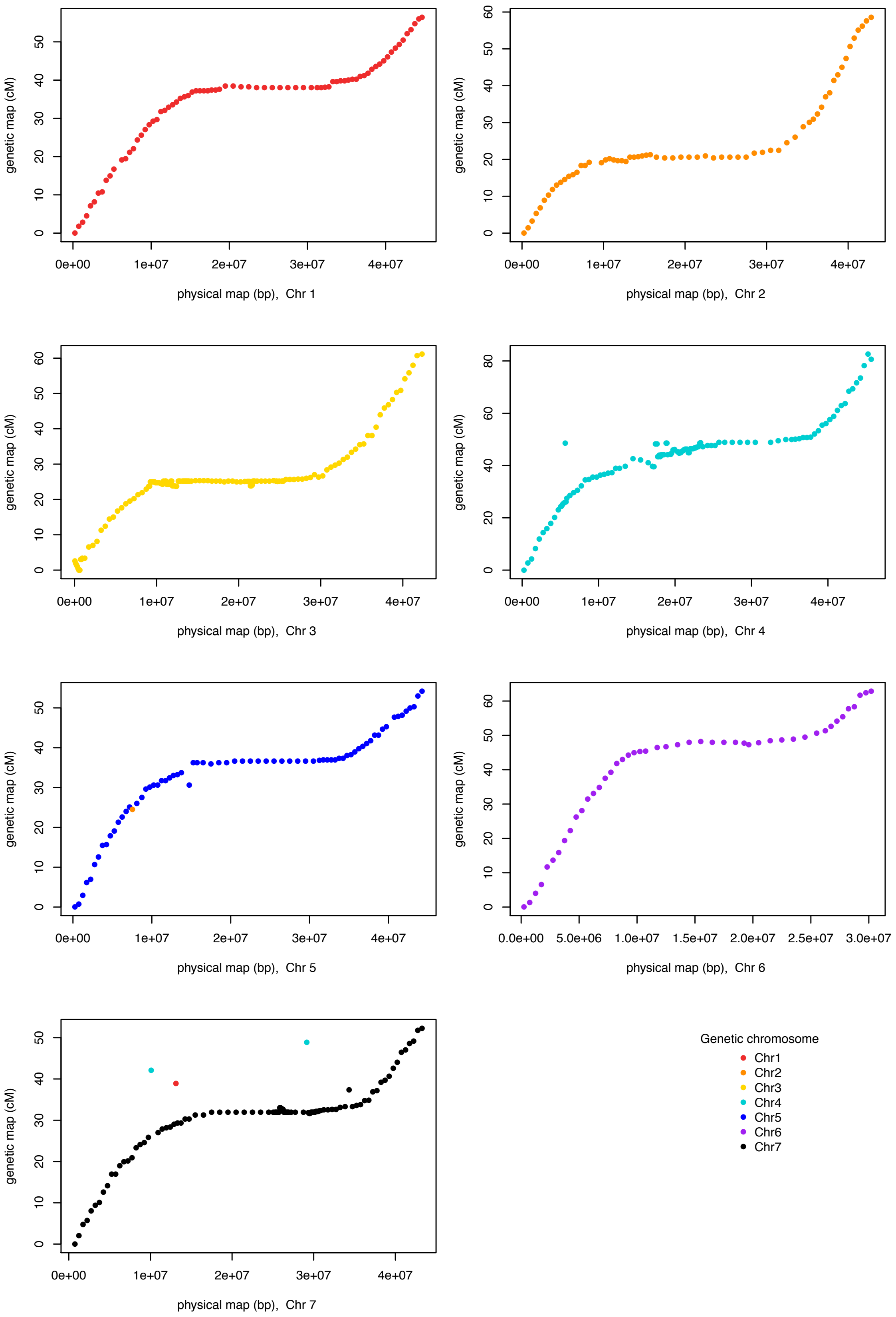

genetic map (cM)
physical map (bp), Chr 1
physical map (bp), Chr 2
physical map (bp), Chr 3
physical map (bp), Chr 4
physical map (bp), Chr 5
physical map (bp), Chr 6
physical map (bp), Chr 7
0e+00
1e+07
2e+07
3e+07
4e+07
0.0e+00
5.0e+06
1.0e+07
1.5e+07
2.0e+07
2.5e+07
3.0e+07
0
10
20
30
40
50
60
80
Genetic chromosome
Chr1
Chr2
Chr3
Chr4
Chr5
Chr6
Chr7

**Figure S3.** Genetic map position versus physical map position of QTL mapping markers. Each box represents the different *Aquilegia* chromosomes (1-7) as assembled in the reference genome (v3.1; e.g., the physical map). The physical map assembly was broken into bins (500 kb or 1 Mb depending on recombination frequency) and bins were genotyped as markers for genetic and QTL mapping. The chromosome that these markers map to in the genetic map are color coded. For the most part, the physical and genetic maps are consistent in marker order and chromosome, however there are several physical map markers that map to different genetic map positions (physical map markers that do not map contiguously in the genetic map) or chromosomes (e.g., several makers on the chromosome 7 physical map to the chromosome 1 or chromosome 4 genetic map). All chromosomes have extended regions of low recombination in the center.

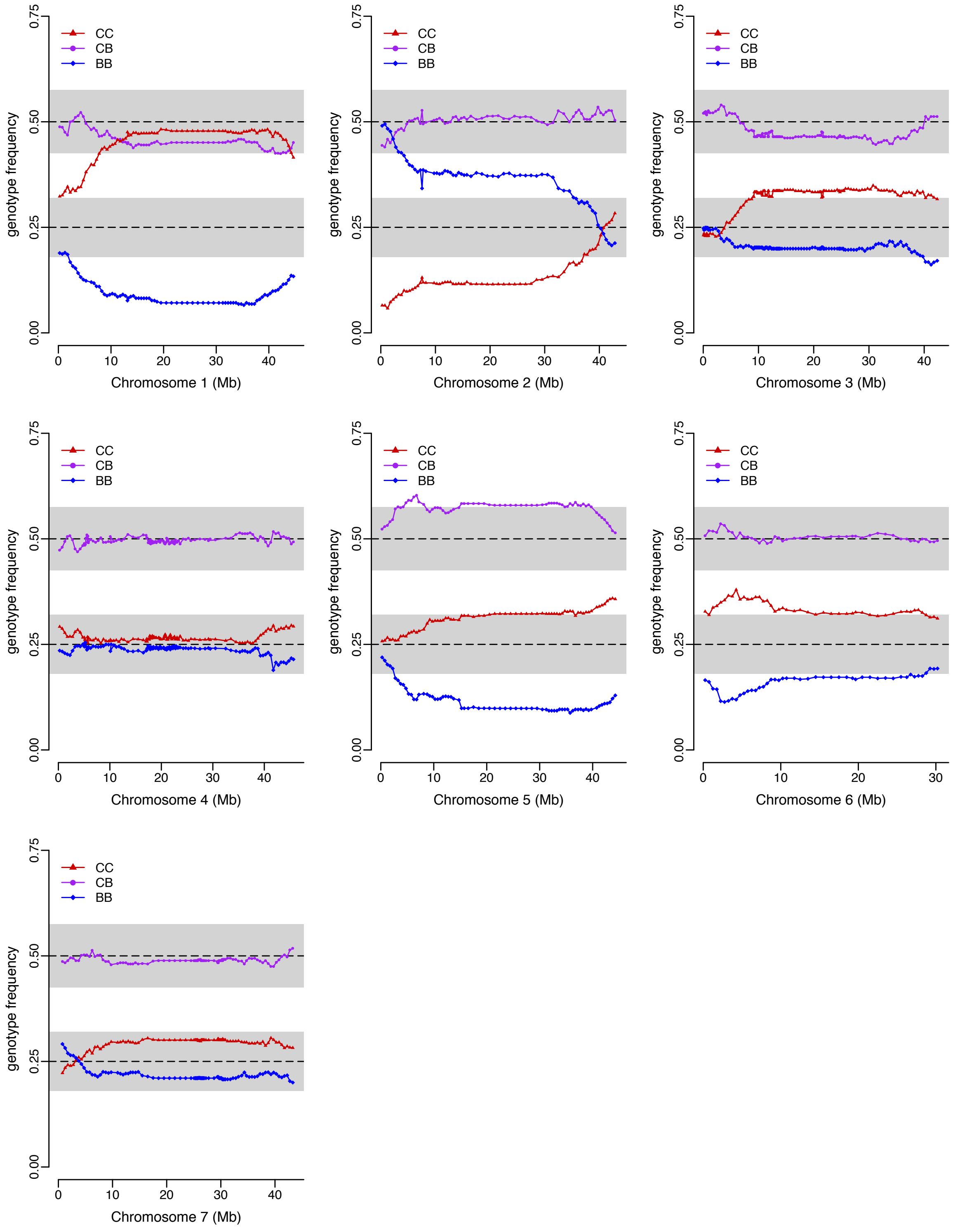


**Figure S4.** F2 population genotype frequency across each chromosome. Loci homozygous for *A. canadensis* alleles are overrepresented across much of chromosomes 1, 3, 5, and 6. Loci homozygous for *A. brevistyla* alleles are overrepresented across much of chromosome 2. CC – homozygous *A. canadensis*, CB – heterozygous, BB – homozygous *A. brevistyla*.

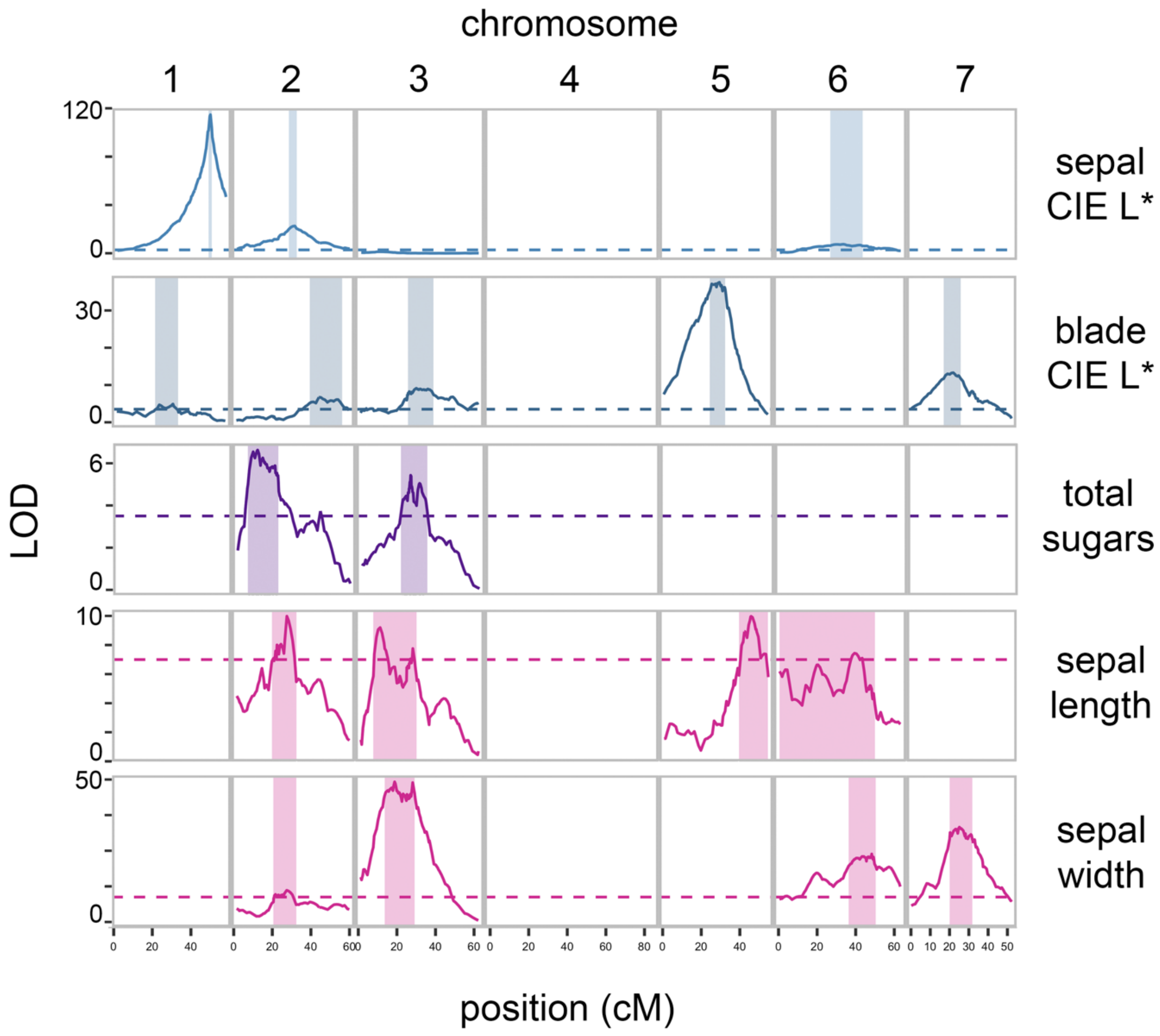


**Figure S5.** Floral trait QTL maps (continued from Fig. 4). Dashed line represents the significant LOD cutoff of 3.5, shaded areas represent the 1.5 LOD interval for each peak.

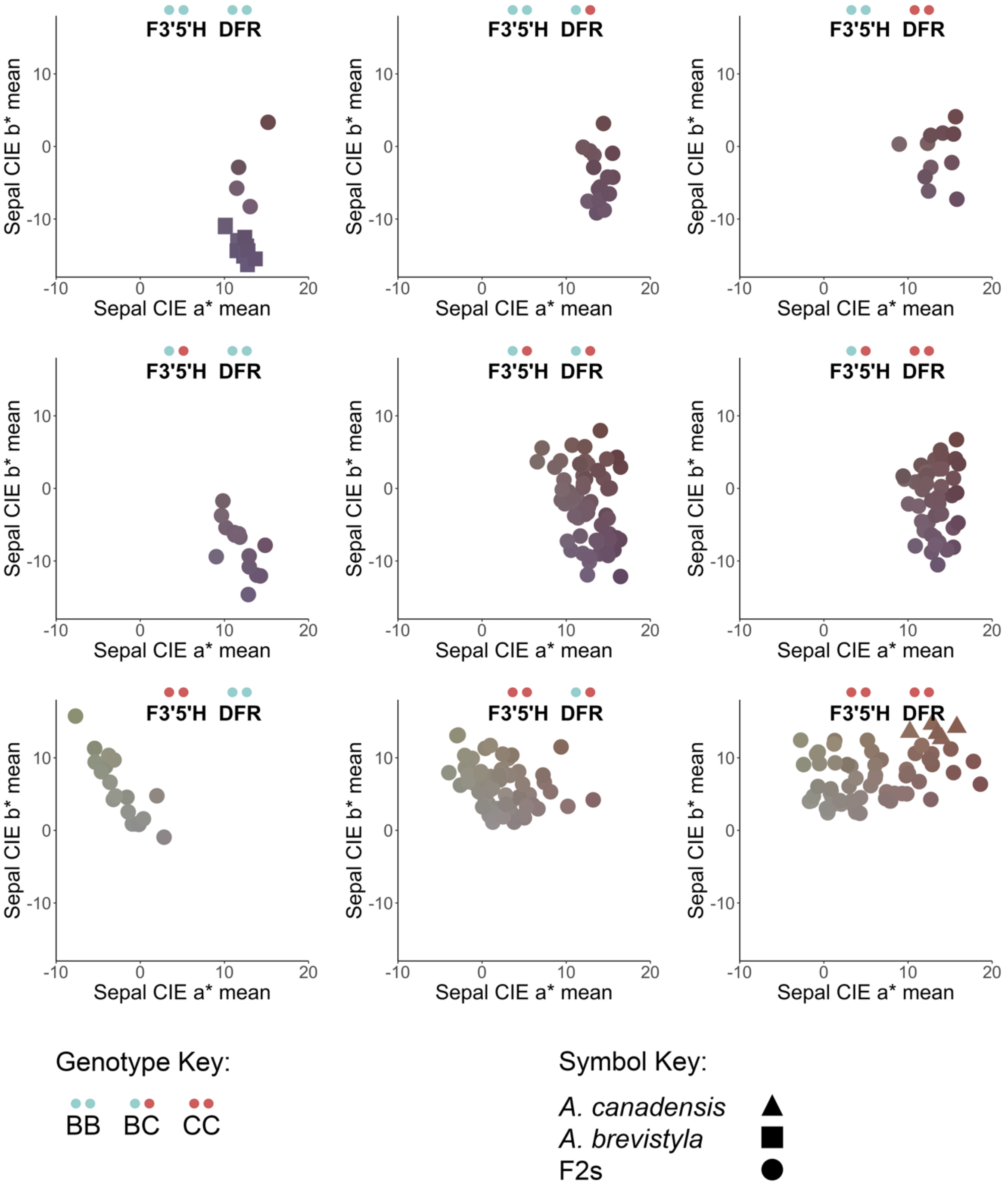


**Figure S6.** Sepal color phenotypes by genotype at *AqF3′5′H* and *AqDFR*. Allelic genotypes of the two loci are represented by pairs of blue (*A. brevistyla* allele) and red (*A. canadensis* allele) dots. Each of the nine panels represents a different allelic genotype combination. Points are color-coded based on the mean RGB values of the sepal for that individual F2 or parent plant. F3′5′H, flavonoid 3′,5′-hydroxylase; DFR, dihydroflavonol reductase.

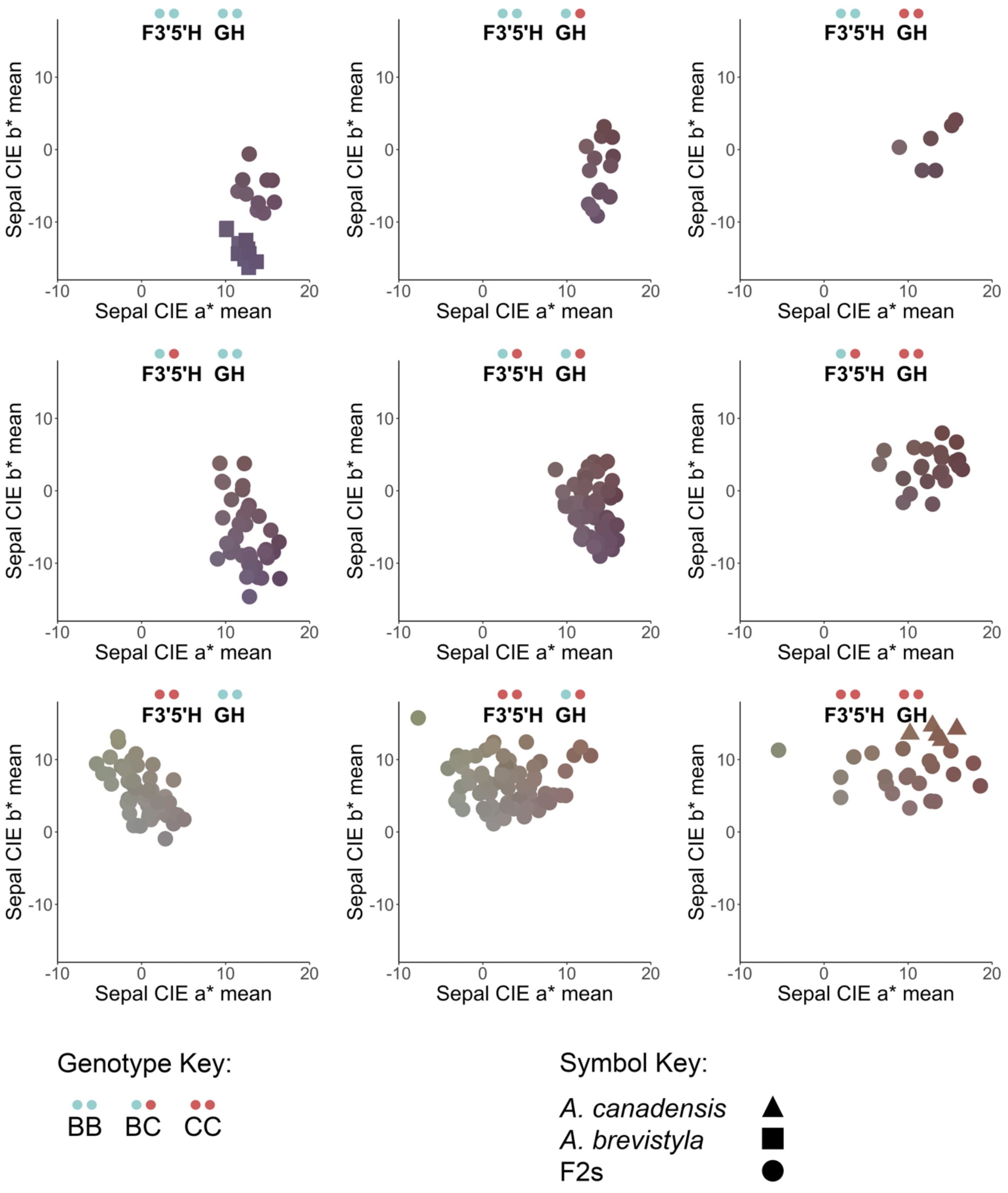


**Figure S7.** Sepal color phenotypes by genotype at *AqF3′5′H* and *AqGH*. Allelic genotypes of each locus are represented by pairs of blue (*A. brevistyla* allele) and red (*A. canadensis* allele) dots. Each of the nine panels represents a different allelic genotype combination. Points are color-coded based on the mean RGB values of the sepal for that individual F2 or parent plant. F3′5′H, flavonoid 3′,5′-hydroxylase; GH, glycosyl hydrolase.

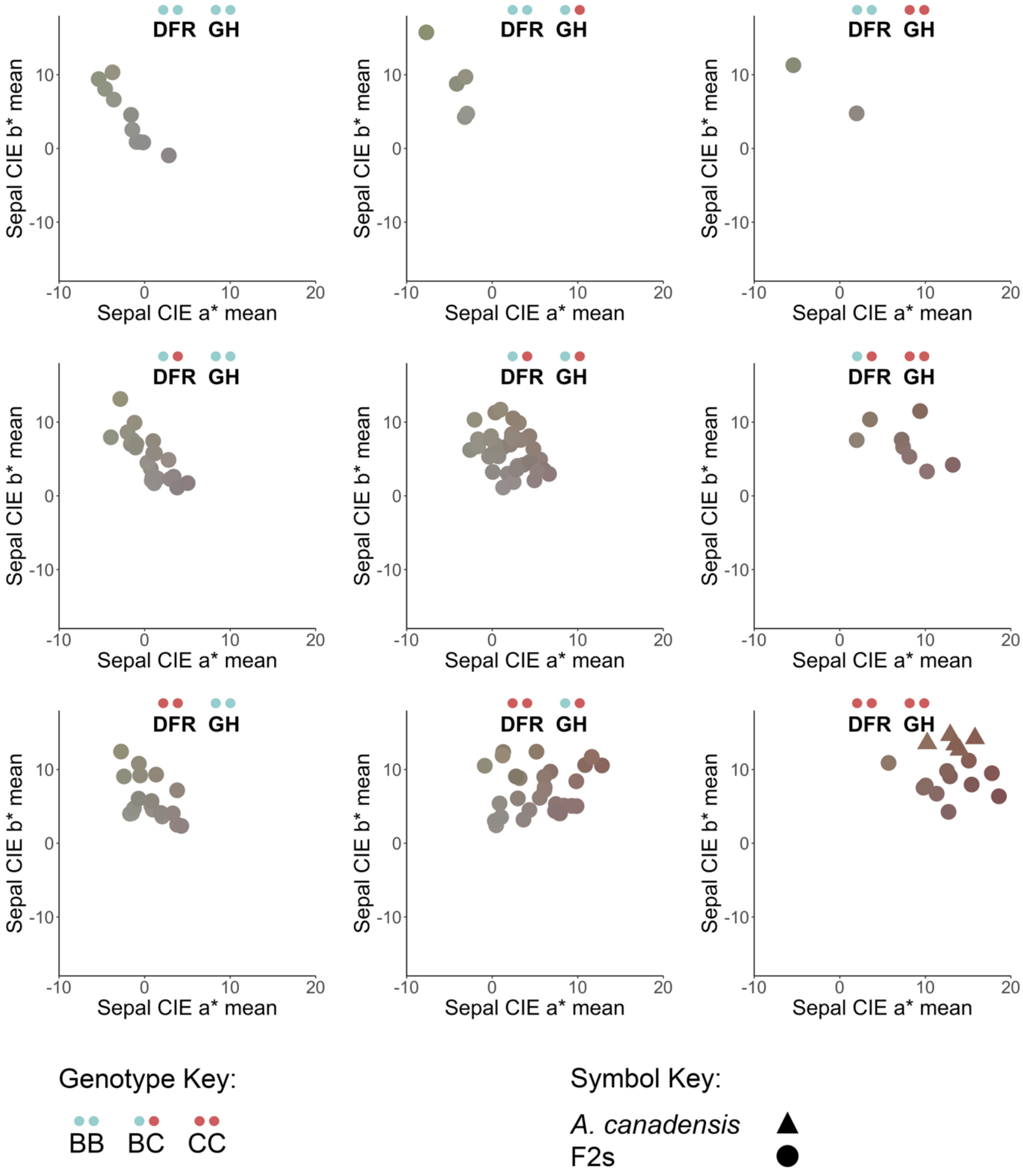


**Figure S8.** Sepal color phenotypes by genotype at *AqDFR* and *AqGH* in plants homozygous for *A. canadensis* at *F3′5′H*. Allelic genotypes of DFR and GH are represented by pairs of blue (*A. brevistyla* allele) and red (*A. canadensis* allele) dots. Each of the nine panels represents a different allelic genotype combination. Points are color-coded based on the mean RGB values of the sepal for that individual F2 or parent plant. F3′5′H, flavonoid 3′,5′-hydroxylase; DFR, dihydroflavonol reductase; GH, glycosyl hydrolase.

| **Traits under study** |
|---|
| Sepal CIE a* |
| Sepal CIE b* |
| Sepal CIE L* |
| Blade CIE a* |
| Blade CIE b* |
| Blade CIE L* |
| Nectar volume |
| Nectar concentration |
| Total sugars |
| Nectary area |
| Sepal area |
| Sepal length |
| Sepal width |
| Blade length |
| Spur length |
| Spur curvature |
| Pistil length |

**Table S1.** List of traits under study.

Han, Yuanji, Miao Wu, Liya Cao, Wangjun Yuan, Meifang Dong, Xiaohui Wang, Weicai Chen, and Fude Shang. 2016. "Characterization of OfWRKY3, a Transcription Factor That Positively Regulates the Carotenoid Cleavage Dioxygenase Gene OfCCD4 in Osmanthus Fragrans." *Plant Mol Biol* 12.

Lloyd, Alan, Austen Brockman, Lyndsey Aguirre, Annabelle Campbell, Alex Bean, Araceli Cantero, and Antonio Gonzalez. 2017. "Advances in the MYB–BHLH–WD Repeat (MBW) Pigment Regulatory Model: Addition of a WRKY Factor and Co-Option of an Anthocyanin MYB for Betalain Regulation." *Plant Cell Physiol.* 11.

Matsuba, Yuki, Nobuhiro Sasaki, Masayuki Tera, Masachika Okamura, Yutaka Abe, Emi Okamoto, Haruka Nakamura, Hisakage Funabashi, Makoto Takatsu, Mikako Saito, Hideaki Matsuoka, Kazuo Nagasawa, and Yoshihiro Ozeki. 2010. "A Novel Glucosylation Reaction on Anthocyanins Catalyzed by Acyl-Glucose–Dependent Glucosyltransferase in the Petals of Carnation and Delphinium." *The Plant Cell* 22(10):3374–89. doi: 10.1105/tpc.110.077487.

Matsui, Kyoko, Yoshimi Umemura, and Masaru Ohme-Takagi. 2008. "AtMYBL2, a Protein with a Single MYB Domain, Acts as a Negative Regulator of Anthocyanin Biosynthesis in Arabidopsis." *The Plant Journal* 55(6):954–67. doi: 10.1111/j.1365-313X.2008.03565.x.

Meng, Yingying, Zuoyi Wang, Yi-Qin Wang, Chongnan Wang, Butuo Zhu, Huan Liu, Wenkai Ji, Jiangqi Wen, Chengcai Chu, Million Tadege, Lifang Niu, and Hao Lin. 2019. “The MYB Activator WHITE PETAL1 Associates with MtTT8 and MtWD40-1 to Regulate Carotenoid-Derived Flower Pigmentation in Medicago Truncatula.” *The Plant Cell* tpc.00480.2019. doi: 10.1105/tpc.19.00480.

Miyahara, Taira, Mariko Takahashi, Yoshihiro Ozeki, and Nobuhiro Sasaki. 2012. “Isolation of an Acyl-Glucose-Dependent Anthocyanin 7-O-Glucosyltransferase from the Monocot Agapanthus Africanus.” *Journal of Plant Physiology* 169(13):1321–26. doi: 10.1016/j.jplph.2012.05.004.

Miyahara, Taira, Tomonori Tani, Mariko Takahashi, Yuzo Nishizaki, Yoshihiro Ozeki, and Nobuhiro Sasaki. 2014. “Isolation of Anthocyanin 7-*O*-Glucosyltransferase from Canterbury Bells (*Campanula Medium*).” *Plant Biotechnology* 31(5):555–59. doi: 10.5511/plantbiotechnology.14.0908a.

Mueller, Lukas A., Christopher D. Goodman, Rebecca A. Silady, and Virginia Walbot. 2000. “AN9, a Petunia Glutathione *S* -Transferase Required for Anthocyanin Sequestration, Is a Flavonoid-Binding Protein.” *Plant Physiology* 123(4):1561–70. doi: 10.1104/pp.123.4.1561.

Nishizaki, Yuzo, Nobuhiro Sasaki, Motoki Yasunaga, Taira Miyahara, Emi Okamoto, Mitsutoshi Okamoto, Yukio Hirose, and Yoshihiro Ozeki. 2014. “Identification of the Glucosyltransferase Gene That Supplies the P-Hydroxybenzoyl-Glucose for 7-Polyacylation of Anthocyanin in Delphinium.” *Journal of Experimental Botany* 65(9):2495–2506. doi: 10.1093/jxb/eru134.

Nishizaki, Yuzo, Motoki Yasunaga, Emi Okamoto, Mitsutoshi Okamoto, Yukio Hirose, Masaatsu Yamaguchi, Yoshihiro Ozeki, and Nobuhiro Sasaki. 2013. “*P* -Hydroxybenzoyl-Glucose Is a Zwitter Donor for the Biosynthesis of 7-Polyacylated Anthocyanin in Delphinium.” *The Plant Cell* 25(10):4150–65. doi: 10.1105/tpc.113.113167.

Ohmiya, Akemi. 2009. “Carotenoid Cleavage Dioxygenases and Their Apocarotenoid Products in Plants.” *Plant Biotechnology* 26(4):351–58. doi: 10.5511/plantbiotechnology.26.351.

Quattrocchio, Francesca, Walter Verweij, Arthur Kroon, Cornelis Spelt, Joseph Mol, and Ronald Koes. 2006. “PH4 of Petunia Is an R2R3 MYB Protein That Activates Vacuolar Acidification through Interactions with Basic-Helix-Loop-Helix Transcription Factors of the Anthocyanin Pathway.” *The Plant Cell* 18(5):1274–91. doi: 10.1105/tpc.105.034041.

Sagawa, Janelle M., Lauren E. Stanley, Amy M. LaFountain, Harry A. Frank, Chang Liu, and Yao-Wu Yuan. 2016. “An R2R3- MYB Transcription Factor Regulates Carotenoid Pigmentation in *Mimulus Lewisii* Flowers.” *New Phytologist* 209(3):1049–57. doi: 10.1111/nph.13647.

Stanley, Lauren E., Baoqing Ding, Wei Sun, Fengjuan Mou, Connor Hill, Shilin Chen, and Yao-Wu Yuan. 2020. “A Tetratricopeptide Repeat Protein Regulates Carotenoid Biosynthesis and Chromoplast Development in Monkeyflowers ( *Mimulus* ).” *The Plant Cell* 32(5):1536–55. doi: 10.1105/tpc.19.00755.

Stanley, Lauren, and Yao-Wu Yuan. 2019. “Transcriptional Regulation of Carotenoid Biosynthesis in Plants: So Many Regulators, So Little Consensus.” *Frontiers in Plant Science* 10:1017. doi: 10.3389/fpls.2019.01017.

Tanaka, Yoshikazu, Nobuhiro Sasaki, and Akemi Ohmiya. 2008. “Biosynthesis of Plant Pigments: Anthocyanins, Betalains and Carotenoids.” *The Plant Journal* 54(4):733–49. doi: 10.1111/j.1365-313X.2008.03447.x.

Verweij, Walter, Cornelis Spelt, Gian-Pietro Di Sansebastiano, Joop Vermeer, Lara Reale, Francesco Ferranti, Ronald Koes, and Francesca Quattrocchio. 2008. “An H+ P-ATPase on the Tonoplast Determines Vacuolar PH and Flower Colour.” *Nature Cell Biology* 10(12):1456–62. doi: 10.1038/ncb1805.

Verweij, Walter, Cornelis E. Spelt, Mattijs Bliek, Michel de Vries, Niek Wit, Marianna Faraco, Ronald Koes, and Francesca M. Quattrocchio. 2016. “Functionally Similar WRKY Proteins Regulate Vacuolar Acidification in Petunia and Hair Development in Arabidopsis.” *The Plant Cell* 28(3):786–803. doi: 10.1105/tpc.15.00608.

Winkel-Shirley, Brenda. 2001. “Flavonoid Biosynthesis. A Colorful Model for Genetics, Biochemistry, Cell Biology, and Biotechnology.” *Plant Physiology* 126(2):485–93. doi: 10.1104/pp.126.2.485.

Yuan, Hui, Junxiang Zhang, Divyashree Nageswaran, and Li Li. 2015. “Carotenoid Metabolism and Regulation in Horticultural Crops.” *Horticulture Research* 2(1):15036. doi: 10.1038/hortres.2015.36.

Zhao, Jian. 2015. “Flavonoid Transport Mechanisms: How to Go, and with Whom.” *Trends in Plant Science* 20(9):576–85. doi: 10.1016/j.tplants.2015.06.007.

Zhou, Xiangjun, Ralf Welsch, Yong Yang, Daniel Álvarez, Matthias Riediger, Hui Yuan, Tara Fish, Jiping Liu, Theodore W. Thannhauser, and Li Li. 2015. “*Arabidopsis* OR Proteins Are the Major Posttranscriptional Regulators of Phytoene Synthase in Controlling Carotenoid Biosynthesis.” *Proceedings of the National Academy of Sciences* 112(11):3558–63. doi: 10.1073/pnas.1420831112.

Zhu, Hui-Fen, Karen Fitzsimmons, Abha Khandelwal, and Robert G. Kranz. 2009. “CPC, a Single-Repeat R3 MYB, Is a Negative Regulator of Anthocyanin Biosynthesis in Arabidopsis.” *Molecular Plant* 2(4):790–802. doi: 10.1093/mp/ssp030.
